# Global hypo-methylation in a subgroup of glioblastoma enriched for an astrocytic signature is associated with increased invasion and altered immune landscape

**DOI:** 10.1101/2022.02.15.480525

**Authors:** James R Boot, Gabriel Rosser, Daliya Kancheva, Claire Vinel, Yau Mun Lim, Nicola Pomella, Xinyu Zhang, Loredana Guglielmi, Denise Sheer, Michael R. Barnes, Sebastian Brandner, Sven Nelander, Kiavash Movahedi, Silvia Marino

## Abstract

We describe a new molecular subgroup of glioblastoma, the most prevalent malignant adult brain tumour, harbouring a bias towards hypomethylation at defined differentially methylated regions. This epigenetic signature correlates with an enrichment for an astrocytic gene signature, which together with the identification of enriched predicted binding sites of transcription factors known to cause demethylation and to be involved in astrocytic/glial lineage specification, point to a shared ontogeny between this glioblastoma subgroup and astroglial progenitors. At functional level, increased invasiveness, at least in part mediated by SRPX2, and macrophage infiltration characterise this glioblastoma subgroup.

## Introduction

Glioblastoma IDH-wildtype (now renamed glioblastoma), is a highly aggressive brain tumour^1^, with an extremely poor prognosis of 15 months median survival from diagnosis^2^. GBM is also the most prevalent primary adult brain tumour with an annual occurrence of approximately five cases per 100,000 people^1,3^ and a mean diagnosis age of 64^4^.

Part of the difficulty in researching and treating glioblastoma is the heterogeneity of the tumour, both at inter- and intra- tumoral levels. Inter-tumoral heterogeneity reflects differences at the genetic level, with the most striking distinction being between IDH-mutant glioblastoma (now renamed astrocytoma, WHO grade 4) and IDH-wildtype glioblastoma. Inter-tumoral heterogeneity is also illustrated by the classification of glioblastomas into different subgroups, such as the transcription-based subgrouping proposed by the Verhaak group^5^, and the DNA methylation-based subgrouping proposed by Sturm *et al*.^6^ However, seminal studies by multiple groups have established that there is also considerable intra- tumoral heterogeneity in glioblastoma^7–9^. Established transcriptional glioblastoma subgroups profiles (Proneural, Neural, Classical and Mesenchymal) are variably expressed in single cells from the same tumours^7^ and tumour cells are characterised by distinct gene expression signatures and cluster separately from one another^8^. Single cells have also been shown to score highly for multiple gene expression signatures creating hybrid states, which further increases the heterogeneity of the tumour cell populations^8^. Couturier *et al.*^9^ found that tumour cells fell into a spectrum of states, whereby cells at one end of the spectrum highly expressed neuronal genes, whilst cells at the other end of the spectrum up-regulated astrocytic genes. Alternatively, glioblastomas can be stratified into more distinct subgroups (IDH, RTKI, RTKII, Mesenchymal, K27 and G34) considering not only mutations and gene expression profiling but also DNA methylation data, tumour location, age distribution and protein markers^6^. These 6 subgroups can be identified based on DNA methylation subtyping alone^10^ and show less heterogeneity^6^, although some degree of heterogeneity has been reported in a small number of cases^11^. The tumour microenvironment also contributes to glioblastoma heterogeneity and the crosstalk between malignant cells and for example the inflammosome is well characterised^12^ with macrophages known to drive the transition of cancer cells towards a mesenchymal-like state^13^.

Whether glioblastoma heterogeneity and its underlying epigenetic makeup is determined by the cell of origin or is acquired during transformation, is a matter of debate. The putative cell of origin is thought to be a stem/progenitor cell that acquires the first genetic and/or epigenetic alterations that promote the formation of the tumour^2^. The debate over the cell of origin in glioblastoma centres around neural stem cells (NSCs) and lineage committed progenitor cells, such as oligodendrocyte, astrocytic and neuronal precursor cells. NSCs are logically the prime candidate for the cell of origin of glioblastoma, because of their self-renewal potential, differentiation plasticity and similarity in their gene expression with glioblastoma stem cells (here called glioblastoma initiating cells - GICs)^14^. A seminal study by Zheng *et al.*^15^ showed that deletion of both *Trp53* and *Pten* by Cre recombinase under the control of the *GFAP* promoter increased the proliferative rate and self-renewal capability of NSCs, whilst also inhibiting their ability to differentiate into specific neural lineages, leading to transformation into high grade malignant gliomas. Importantly, glioblastoma driver mutations have been identified in NSCs of the human subventricular zone (SVZ) in tissue samples obtained from patients, providing for the first-time evidence that these cells can act as cells of origin of glioblastoma in human^16^. Further studies^17^ have shown that inducing null alleles of *Nf1*, *Trp53* and *Pten* using Cre recombinase under the control of NSC specific *Nestin*, but also oligodendrocyte progenitor cell specific *NG2* or bipotential progenitor cell - specific *Ascl1* led to high grade glioma, whilst no tumours formed when the same null alleles were induced in mature neurons (*Camk2a Cre)*, immature neurons (*Neurod1 Cre)* and adult neuronal progenitors (*Dtx1 Cre)*^18^. These studies together with others which have shown that also OPC^19,20^ and astrocytes^21^ can behave as cell of origin of IDH-wildtype glioblastoma raise the possibility that different subgroups may originate from glial progenitor cells at different developmental stages.

Here, we leveraged pairs of patient-derived GICs and patient-matched expanded potential stem cells (EPSC) - derived neural stem and progenitor cells to investigate the DNA methylation landscape of GICs as compared to their putative cell of origin in a patient-specific manner to further characterise glioblastoma heterogeneity and its ontogeny.

## Materials and Methods

### Tissue Culture

We have previously described a novel experimental pipeline, SYNGN, to derive GICs and EPSCs as well as induced NSC (iNSCs) from patients who underwent surgical resection of glioblastoma^22^. The use of human tissue samples was licensed by the National Research Ethics Service (NRES), University College London Hospitals NRES license for using human tissue samples: Project ref 08/0077 (S. Brandner); Amendment 1 17/10/2014. In brief, at the time of operation, tumour tissue and a thin strip of the dura mater were obtained. Fibroblasts were isolated from the dura mater, propagated, and reprogrammed to generate EPSCs, which were further differentiated into iNSCs, induced Astrocyte Progenitors (iAPCs) and induced Oligodendrocyte Progenitors (iOPCs). All media recipes can be found in Supplementary Tables 1-5.

GICs were cultured on laminin (Sigma Cat. #L2020) coated tissue culture plates at 37°C, 5% CO_2_ . Cells were maintained in Neurocult media (Stem Cell Technologies Cat. #05751) supplemented with 1% penicillin/streptomycin solution (Sigma Cat. #P4458), Heparin (Stem Cell Technologies Cat. #07980), EGF (Peprotech Cat. #AF-315-09-1MG) and FGF (Peprotech Cat. #AF-100-18B-50UG) and passaged once they reached 80-90% confluence. GICs from the HGCC cohort were cultured on Poly-L-Ornithine (Sigma Cat. #P3655) and laminin coated plates and maintained in media termed HGCC Media (Supplementary Table 1), supplemented with EGF and FGF, and passaged in the same manner as GICs from our own cohort. Separate passages of the same GIC line were considered to be biological replicates.

Dura-derived fibroblasts were reprogrammed into EPSCs as previously published^23,24^, EPSCs were then further differentiated into iNSCs with a Gibco commercially available kit^22^ (Gibco Cat. #A1647801). iNSCs were cultured on GelTrex (Gibco Cat. #A1413302) coated tissue culture plates and maintained in Neural Expansion media (Supplementary Table 2) at 37°C, 5% CO_2_ . iNSCs were passaged once they reached 80-90% confluence.

HEK293T cells were cultured in adherent conditions and maintained in IMDM media (Gibco Cat. #12440061) supplemented with 10% Foetal Bovine Serum (FBS) (Gibco Cat. #10500064) and 1% penicillin/streptomycin solution (Sigma Cat. #P4458) at 37°C, 5% CO_2_ . They were detached using 1X Accutase (Millipore Cat. SCR005) for five minutes and passaged at ratios from 1:5 to 1:50.

### iAPC and iOPC Generation

Differentiation of iNSCs into iAPCs was adapted from published protocols^25^. Differentiation was initiated on Day -1, by seeding dissociated iNSCs at 1.5×10^4^ cells/cm^2^ density on GelTrex-coated plates in Neural Expansion media. On Day 0 Neural Expansion media was changed to Astrocyte media (ScienCell: Cat. #1801) supplemented with 2% FBS, Astrocyte Growth Supplement (AGS) and penicillin/streptomycin solution, provided with the media. From Day 2 onwards, media was changed every 48 hours for 20 – 30 days. At 80-90% confluence, the cells were passaged back to the starting seeding density (1.5×10^4^ cells/cm^2^), or at an approximate ratio of 1:6. Cells were detached using Accutase, and always cultured in the same Astrocyte media on GelTrex. Cell pellets were collected at three different time points throughout the differentiation process, when cells were confluent and passaged; approximately at days 10, 20 and 30. Differentiated astrocytes could be cryopreserved using Astrocyte media supplemented with 10% DMSO, or commercially available SynthFreeze (Gibco, Cat. #A1254201).

iNSCs were differentiated into iOPCs using a published protocol^26^, commencing from iPSCs and achieving fully mature oligodendrocytes at 95-days. Here, the protocol was started from iNSCs, equivalent to Day 8 of the Douvaras-Fossati protocol, which were induced into OPCs up to Day 75, when the original authors reported emergence of immature O4^+^ oligodendrocytes. On Day 0 of our protocol (Day 8 of the Douvaras-Fossati protocol) Neural Expansion media was removed, and iNSCs cultured in N2 media (Supplementary Table 3) with 100 nM RA and 1 µM Smoothened Agonist (SAG) added freshly each day. Media was then changed daily with fresh RA and SAG until Day 4, at which point cells became over-confluent and were detached and placed in N2B27 Media (Supplementary Table 4) with freshly added RA and SAG. A series of scratches were made with a cell scraper vertically, horizontally, and diagonally across each well, and the contents of a single well transferred into eight wells of an ultra-low attachment 24-well plate (Corning Cat. #3473), with extra N2B27 media. Aggregates were then cultured in suspension for a further eight days, with media changed every 48 hours. On Day 12 of the protocol N2B27 media was replaced with PDGF media (Supplementary Table 5), and aggregates were cultured for a further 10 days, with media replaced every 48 hours. On Day 22 of the protocol aggregates were plated onto Poly-L-Ornithine and laminin coated plates and were cultured adherently until Day 67 of the protocol, with PDGF media changes every 48 hours.

Two independent differentiations of iAPCs and iOPCs from iNSCs were considered to be biological replicates. For iNSCs two independent differentiations from iPSCs were considered to be biological replicates.

### Proliferation assay

For the comparison of iNSCs and iAPCs cells were seeded at 2×10^4^ cells per well of a 24-well plate (CytoOne Cat. #CC7682-7524), whilst for the comparison of GIC lines cells were seeded at 1×10^4^ cells per well of a 24-well plate. Then, at selected time-points (every 24 hours) starting from Day 1 or Day 2, cells from each well were individually detached using Accutase, then centrifuged individually (1200 rpm) at 4°C for five minutes. At least three wells were detached and counted at each timepoint to generate technical triplicates. After detachment and centrifugation, cell pellets were resuspended in 100 µL DPBS and 10 µL was mixed with 10 µL of Trypan Blue (Sigma Cat. T8154-100ML) and live cells counted using a haemocytometer.

### Invasion assay

Transwell inserts with 8.0 µm pores (Sarstedt Cat. #89.3932.800) were placed into wells of a 24-well plate and coated with 100 µL of GelTrex. A total of 100,000 cells were then seeded into the transwell insert in 200 µL and 700 µL of normal growth media was added to the bottom of the well. Cells were then incubated in normal growth conditions for 24 hours, at which point cells on the inside of the transwell were removed using a cotton bud dampened with DPBS. Once cells inside the transwell were removed, cells on the bottom of the transwell were fixed using methanol, pre-chilled to -20°C, for five minutes at room temperature. After fixation, the bottom of the transwell was washed twice for five minutes using DPBS. The membrane of the transwell was then cut out and mounted onto a microscopy slide with mounting media including DAPI (Vectorlabs Cat. #H-2000). Transwell membranes were then analysed using a microscope and five representative images of nuclei on each membrane captured. For each biological group and replicate (different passages of cell lines), three technical replicate membranes were imaged. Finally, the number of whole nuclei in each image field were counted, using ImageJ software (Version 1.51m9), to ascertain how many cells migrated across the membrane.

### Neurosphere extreme limiting dilution assay

On Day 0 of the assay, cells were seeded at a maximum cell density of 25 cells per well of a round bottom ultra-low attachment 96-well plate (Corning Cat. #7007). Cells at this density were then serially diluted 1:2 a total of 4 times to give 5 cell densities in total – 25, 12.5, 6.25, 3.125 and 1.5625 cells per well. For each biological replicate (different passages of cell lines), 12 technical replicates of each cell density were performed (12 separate wells). Cells were then incubated for 14-days with media changes every 48 hours, after which time the presence of neurospheres was assessed and counted for each well. To be considered a neurosphere, cells had to form 3D spherical clusters with smooth and defined edges and had to be greater than two cells in size. Results were analysed using the extreme limiting dilution analysis tool ^27^, where the log proportion of negative cultures is plotted against the number of cells seeded, with a trend line indicating the estimated active stem cell frequency. The statistical significance of the differences between the estimated active stem cell frequencies of different cell lines was also tested using a Chi Square test as part of the analysis tool ^27^.

### Animal procedures

All procedures were performed in accordance with licenses held under the UK Animals (Home Office Guidelines: animals Scientific Procedures Act 1986, PPL 70/6452 and P78B6C064Scientific Procedures) Act 1986 and later modifications and conforming to all relevant guidelines and regulations. Orthotopic xenografts were performed on eight-to twelve-week-old NOD SCID CB17-Prkdcscid/J mice (purchased from The Jackson laboratory) under anaesthesia with isoflurane gas and 5 × 10^5^ primary human GIC in 10 µL PBS were slowly injected with a 26-gauge Hamilton syringe needle into the right cerebral hemisphere with the following coordinates from the bregma suture: 2 mm posterior, 2 mm lateral, 4 mm deep, 10° angle. After the injection, scalps were cleaned with ethanol swab to remove any remaining cells and sutured with 4-0 Coated Vicryl Suture (Ethicon). After the surgery, mice recovered on a heat-map until they were fully awake. For the five days following surgery, mice were checked twice a day, then once a day and body weight was monitored once a week. Mice were kept on tumour watch until they developed brain tumour clinical symptoms and were then euthanized by neck dislocation and brains were harvested for histology analysis.

### DNA and RNA extraction

Cells used for qPCR analysis were pelleted and frozen at -80°C before RNA extraction. RNA was extracted by following the standard protocols of either Qiagen RNeasy Mini or Micro kits (Cat. #74104/74004). Some cell pellets were processed using Norgen Biotek RNA/DNA/Protein Purification Plus Kit (Cat. #47700), which allows genomic DNA, total RNA, and protein to be isolated from a single sample. RNA to be sent for RNAseq, and DNA to be sent for DNA-Methylation array, were prepared using the Norgen Biotek kit, according to manufacturer’s instructions.

### Reverse Transcription and qPCR

Reverse transcription reactions were carried out by first mixing 1 µL random primers (Invitrogen Cat. #48190011), 500 ng RNA and ddH_2_ O up to 10 µL, primers were then annealed by heating to 65°C for five minutes, then 4°C for five minutes. Then 4 µL 5X FS Buffer, 1 µL 0.1 M DTT, 0.5 µL SuperScript III Reverse Transcriptase (Invitrogen Cat. #18080044), 1 µL 10 mM dNTP mix (Invitrogen Cat. #18427-013) and 3.5 µL ddH_2_O were added to this reaction mixture.

qPCR reactions were carried out using Applied Biosystems Syber Green qPCR Master Mix (Cat. #4309155). Each reaction contained 2 µL of 2.5 ng/µL cDNA (5 ng total), 0.48 µL of 10 µM forward and reverse primer mix, 3.52 µL ddH_2_ O and 6 µL of Syber Green Master Mix (12 µL total reaction volume). All qPCR reactions were run on an Applied Biosystems 7500 Real-Time PCR System or StepOnePlus™ Real-Time PCR System. A full list of primers and their sequences used throughout this project can be found in the Supplementary Table 6.

### Flow cytometry and Fluorescent Activated Cell Sorting (FACS)

Primary and secondary antibodies used for flow cytometry and FACS staining are listed in Supplementary Table 7. Samples stained with unconjugated primary antibodies were incubated with species reactive secondary antibodies with various fluorophores conjugated. Samples for flow cytometry analysis were analysed using a BD LSRII Analyser, samples for FACS were processed using a BD FACS Aria Sorter.

For FACS of live iOPCs, cells were dissociated by removing culture media, adding FACS buffer (1:200 BSA Sigma Cat. #A3912, 1:250 EDTA 0.5mM Ambion Cat. #AM9262, in DPBS) and homogenising the cells into a single cell suspension. As the culture contained large aggregates, the cell suspension was passed through a 100 µm size filter. Cells were then counted and centrifuged at 1500 rpm for five minutes at 4°C followed by dispensing 100 µL of a 5×10^6^ cells/mL suspension per tube for staining. Prior to staining cells were incubated in anti-CD16/32 FcR block (diluted 1:200 in FACS buffer) for 15 minutes at 4°C, washed and then incubated with conjugated or unconjugated primary antibodies for 30 minutes at 4°C. Staining was carried out with either single antibodies or combinations of antibodies. Finally, cells were pelleted and re-suspended in DPBS before FACS.

For flow cytometry analysis of cell samples, cells were harvested, resuspended in FACS buffer, and blocked with FcR blocker in the same manner as for FACS analysis. Extracellular antibodies were first incubated with samples for 30 minutes at 4°C, followed by washing and resuspension in a fixable viability dye diluted 1:200 in DPBS and incubated for 20 minutes at 4°C. After further washing, cells were then fixed for 20 minutes at 4°C in 4% PFA diluted in a 1:1 ratio with FACS buffer. For the staining of intracellular targets, after fixation the cells were permeabilised using methanol for five minutes at room temperature and incubated with antibodies diluted in methanol for 20 minutes at room temperature. Finally, samples were washed in FACS buffer and resuspended for analysis.

### Immunocytochemistry (ICC)

All cells analysed via immunocytochemistry (ICC) were washed in DPBS and fixed by treatment with 4% paraformaldehyde for 15 minutes, then washed in DPBS for five minutes, three times. Fixed cells were stored at 4°C in DPBS until staining. Cells were then permeabilised and blocked followed by staining with primary antibodies. Primary antibodies and the dilutions used in this study can be found in the Supplementary Table 7. After primary antibody incubation, overnight at 4°C or three hours at room temperature, samples were washed in DPBS, and stained with species reactive secondary antibodies conjugated to various fluorophores for one hour at room temperature. After washing once again in DPBS, sample slides were then mounted using Fluoroshield mounting media with DAPI and sealed using nail varnish.

### Immunoblotting

Cell samples used for protein extraction were first pelleted by centrifugation, at full speed, for five minutes, then snap frozen using dry ice and stored at -20°C until extraction. Protein was extracted from cell pellets by resuspending pellets in RIPA lysis buffer (25 mM Tris-HCl pH 7.6, 150 mM NaCl, 1% NP- 40, 1% sodium deoxycholate, 0.1% SDS) supplemented with protease inhibitors (Santa Cruz Cat. SC- 24948A), samples were then left to incubate on ice for 30 minutes. After incubation samples were centrifuged, at full speed, for 15 minutes, at 4°C, and the supernatant collected.

Protein concentration was determined using BCA assay (Thermo Cat. #23227) performed as per the manufacturer’s instructions. Concentration of protein samples was then determined by interpolating the absorbance values of the unknown samples with a standard curve of known protein concentrations.

An equal amount of protein was then loaded into a 4-12% acrylamide gel (Invitrogen Cat. #NP0322BOX). Proteins were separated in SDS-PAGE (ThermoFisher Cat. #NP-0001) and blotted onto a nitrocellulose membrane (GE Healthcare Cat. #10600002). Membranes were then blocked with 5% non-fat milk (Santa Cruz Cat. #SC-2325) in Phosphate Buffered Saline-Tween (PBS-T) (0.1% Tween20 (Sigma Cat. #P9416-100ML) in PBS) for one hour at room temperature and then incubated with primary antibodies at 4°C overnight. The Supplementary Table 7 summarises primary antibodies and their dilutions used in this study. After incubation of primary antibody membranes were washed three times for five minutes in PBS-T and then incubated, at room temperature, with the species appropriate peroxidase-conjugated secondary antibody at a dilution of 1:5000 for one hour. Membranes were further washed three times for five minutes in PBS-T before being visualised using ECL kit (GE Healthcare Cat. #RPN2232) and ChemiDoc Imaging System (BioRad). Quantification of the protein expression was measured by densitometric analysis performed with ImageJ software (Version 1.51m9).

### Enzyme linked immunosorbent assay (ELISA)

Quantification of secreted proteins and chemokines such as Interleukin-6 (IL-6) was performed using an ELISA. Cells were cultured, prepared, and treated as already described, and after treatment growth media supernatant was collected for analysis by ELISA. Upon collection, supernatant was filtered using 0.22 µm syringe filter (Santa Cruz Cat. #SC-358812) and snap frozen using dry ice, followed by storage at -80°C until analysis. The ELISA was then performed as per the manufacturer’s instructions (BD Bioscience IL-6 ELISA Cat. #555220), and concentrations determined by interpolating absorbance values of samples using a standard curve.

### shRNA lentivirus production and transduction of cell lines

Lentivirus particles were produced using HEK293T cells. To produce short hairpin RNA (shRNA) lentivirus’, a Lipofectamine3000 (Invitrogen Cat. #L3000-015) transfection protocol was used. Transfection was carried out as per the manufacturer’s standard protocol, by forming DNA-lipid complexes which were then incubated on cells for six hours followed by addition of packaging media for lentivirus harvesting. Packaging media consisted of Optimem reduced serum media supplemented with Glutamax (Gibco Cat. #51985-034), with 1 mM sodium pyruvate (Gibco Cat. #11360-039) and 5% FBS. 24 hours post transfection, media was harvested from HEK293T cells and stored at 4°C, 10 mL of fresh packaging media was added. 52 hours post transfection media was harvested once again and mixed with media previously harvested. Harvested supernatant was then centrifuged at 2000 rpm for five minutes to remove cell debris, then filtered through a 0.44 µm filter (VWR Cat. #514-0329). To precipitate and concentrate lentiviral particles, 5X Polyethylene glycol (PEG) (Sigma #89510-1KG-F) was prepared by mixing 200g PEG, 12g NaCl (Fisher Cat. #S/3166/60) and 1 mL of 1M Tris (pH 7.5) (PanReac AppliChem Cat #A4263,0500) then ddH_2_ O added to a total volume of 500 mL. pH of PEG was further adjusted to 7.2, and then autoclaved before use. 5X PEG was then added to harvested supernatant to give a final concentration of 1X and then incubated at 4°C overnight. The following day harvested supernatant mixed with PEG was centrifuged at 1500g for 30 minutes at 4°C, supernatant was removed and spun again to remove excess supernatant. The lentiviral pellet was then resuspended in an appropriate amount of DPBS, aliquoted and stored at -80°C until use.

Harvested virus was titrated to determine the transforming units per mL (TU/mL) for each volume of virus used during titration. Cell lines were infected overnight with the stated multiplicity of infection (MOI) of lentiviral particles. The TU/mL that achieved 5% to 30% of fluorescent tag positive cells during lentivirus titration was selected for the calculation of the MOI. The following day after infection with lentivirus, media was changed, and the cells were left to recover. Once confluent the cells expressing the desired construct were then purified by Puromycin selection or FACS, if transduced with constructs with a Puromycin resistance gene or fluorescent tag.

Glycerol stocks of competent bacteria containing shRNA plasmids, for lentivirus production, were purchased from Horizon Discovery Dharmacon™. To obtain plasmid for lentiviral production, a small of amount of glycerol stock was extracted and added to LB broth (Sigma Cat. #L3522) supplemented with the relevant antibiotic, and grown up in large liquid culture, overnight at 37°C. Following this, plasmid DNA was isolated and purified using the Qiagen Maxi Prep kit (Qiagen Cat. #12963). Details of each shRNA plasmid constructs can be found in the Supplementary Table 8.

### DNA methylation data processing

DNA used for methylation arrays was extracted and prepared as described above. Two biological replicates of each patient-matched GICs, iNSCs, iAPCs, and iOPCs were sent for DNA methylation array. DNA was assayed on the Illumina Infinium Methylation EPIC array (over 850,000 probes). Raw data was imported into an R workspace (R Version 3.5.0) and all analysis therein performed using the RStudio environment (Version 1.1.453). Raw data from the array was first processed using the ChAMP^28,29^ (Version 2.12.4) R package to remove any failed detections and flawed probes. Along with data, metadata was also imported and used to help perform analysis. After this initial processing, data were then further processed and normalised using the Subset-quantile within array normalization (SWAN) algorithm^30^. All methylation data used for further analysis such as differential methylation, PCA or heatmap and dendrograms were SWAN normalised data.

After initial processing of DNA methylation array data, as described above, differentially methylated regions (DMRs) and genes were identified. To do so, output data from the SWAN normalisation algorithm in the form of beta values were imported into an R workspace and different datasets merged into a single dataset with the only probes that were common to all samples. After initial data processing, the R package DMRcate^31^ (Version 1.18.0) was then used to first identify DMRs and then the corresponding genes. The minimum number of contiguous (or consecutive) differentially methylated probes per region to be called a differentially methylated region (DMR) was set to 6, the beta cut off was set to 0.3, and as the lambda parameter (bandwidth) was set to 1000, the scaling factor (C) was set to 2 as per the authors recommendations. Unless otherwise stated here, all other parameters were set to their respective default settings.

### RNAseq data processing

RNA used for RNAseq was extracted and prepared as previously detailed. RNA samples from two biological replicates of patient-matched iAPCs, iNSCs and GICs was of sufficient quality and quantity (minimum 1000 ng total mass at concentration of 50 ng/µL with 260:280 and 260:230 ratios equal to ∼2) as to perform Poly-A library preparation followed by sequencing using the Illumina HiSeq4000 platform at 75 PE. For sequencing of miRNAs, total RNA extracted as already outlined was prepared using the NEBNext smallRNA kit for Illumina (E7330L). In some cases where insufficient quantities of RNA were isolated from single passages of cells, RNA from 2 passages were combined. iOPC samples yielded very low quality and quantity of RNA after extraction and therefore samples were not suitable to be prepared using the Poly-A library preparation method. Instead, two biological replicate iOPC samples, for each patient, were prepared using the SmartSeq2 library preparation method and sequenced using the Illumina HiSeq4000 platform at 75 PE.

The raw RNAseq data generated was processed in multiple ways depending on the output of the analysis. FastQC (v. 0.11.5) (https://www.bioinformatics.babraham.ac.uk/projects/fastqc/) was used to perform quality control of the raw data, to check the Phred score, the GC content distribution, and the duplication levels, then the TrimGalore tool (v. 0.4.5-1) (DOI 10.5281/zenodo.5127898) was used to remove low quality reads (Phred score < 20) and residual adapters. For the use of differential expression analysis raw data was processed using the STAR gapped alignment software^32^ (Version 2.7.0), which generated gene counts. The reference genome used for alignment was Ensembl GRCh38 (release 90). Following alignment, prior to differential gene expression analysis, further processing of the STAR output data was performed. Genes with a counts per million (CPM) value < 1, in a minimum of half of the samples per group (i.e., GIC or iNSC) plus one, were removed. DEGs were identified using the R package: EdgeR^33,34^ (Version 3.24.3) with the thresholds that the minimum absolute log fold change (logFC) in gene expression was 2 and the false discovery rate (FDR) was less than 0.01. EdgeR analysis was performed using two statistical tests provided by the package – the likelihood ratio test (glmFit) and the quasi-likelihood ratio test (glmQLFit), however all gene lists used in further downstream analysis were generated from the more conservative glmQLFit test, unless otherwise stated.

For the differential gene expression analysis of GICs from the HGCC cohort, microarray data that had been pre-processed (normalised and Combat batch adjusted) was used. Differential gene expression analysis was then performed in an R workspace using the RStudio environment and limma package^35^ (Version 3.40.6). Linear modelling was implemented by the lmFit function and the empirical Bayes statistics implemented by the eBayes function and DEGs with a p-value < 0.05, and log fold change > 0.5, were selected by the topTable function. Volcano plots depicting the LFC and statistical significance of DEGs, from analysis of both microarray data and RNAseq data, were generated using the R package EnhancedVolcano (version 1.5.10).

To generate heatmaps, dendrograms and perform PCA (all detailed below), a different approach was used to process raw RNAseq data. Alignment was instead performed using pseudoalignment package Salmon^36^ (Version 0.13.1), with the output being transcript expression level, which were then pooled to give gene level expression estimates expressed as transcripts per million (TPM)^37,38^. This unit is normalised for library size and transcript length. The reference genome used for Salmon alignment was Ensembl GRCh38 (release 90).

### Principal component analysis (PCA), heatmaps and dendrograms

PCA was performed on both methylation array data and RNAseq data to validate the sequencing data. Output data from the pseudoalignment package Salmon (TPM counts) were used for PCA of RNAseq data, and beta values from SWAN normalised data we used for PCA of methylation array data. Prior to PCA, data were filtered to include only the most variable methylation probes or genes, the exact number used for each figure is stated in the relevant figure legend. The R package NOIseq^39,40^ (Version 2.26.1) was used to perform PCA and then the results plotted, using R packages ggplot2^41^ (Version 3.1.1) (for two-dimensional plotting of two principal components) or plotly (Version 4.9.0) (for three-dimensional plotting of three principal components).

Heatmaps and dendrograms were generated using both DNA methylation and RNAseq data. The RNAseq data used was the output from the pseudoalignment package Salmon (gene expression data in units of TPM). DNA methylation data used was beta values from SWAN normalisation. RNAseq data was filtered prior to analysis by removing genes with a TPM value < 1 and the number of samples in each group (i.e., GIC or iNSC) that must express each gene to half the group size plus one. After this initial filtering, only the most variable genes were used in the analysis, with the exact number stated for each figure. In general, the Euclidean clustering method and Complete distance methods were used as part of the standard heatmap.2 function in R.

### Gene signature analysis

Gene signature analysis was performed using Single Sample Gene Set Enrichment Analysis (ssGSEA) method. In order to calculate ssGSEA enrichment scores for our samples, the R package GSVA^42^ (Version 1.30.0) was used. The gene level expression estimates output, expressed as TPM, from alignment using the Salmon (detailed earlier) were used as the input for this analysis. As well as gene expression data, gene lists or signatures were also required, and all gene lists/signatures were formatted as Ensembl gene IDs.

The oligodendrocyte gene signatures used here were taken from a published study by McKenzie *et al.*^43^ The astrocyte composite signature (ACS) was manually generated by compiling multiple astrocytic gene signatures into one coherent gene signature. Astrocytic gene signatures were first found in the xCell bioinformatics tool^44^: xCell is a tool used to de-convolute a sample composed of a mixture of cell types into its respective cell types, based on gene signatures curated and validated by the authors. Multiple data sources were used to generate gene signatures for as many cell types as possible, and the authors generated a gene signature for a given cell type from each data source, meaning more than one gene signature was generated for each cell type. For example: three data sources contained astrocyte expression data, and thus two astrocyte signatures were generated from each data source, meaning there were six signatures for the tool to use to de-convolute mixed samples. In this present study, we have taken these six astrocyte signatures used by the xCell tool and merged them into one coherent signature. The ACS and the oligodendrocyte signatures used in this study can be found in Supplementary Table 9.

### Motif Analysis using Homer

Identification of enriched binding motifs in genomic regions was performed using Homer^45^ (Hypergeometric Optimization of Motif EnRichment, (v4.11, 10-24-2019). The tool findMotifsGenome.pl was used to perform *de novo* search as well as to check the enrichment of known motifs, in the context of the latest human genome annotation (hg38). Homer searched significantly enriched motifs (p-value < 0.05) with a length spanning a wide range of standard values (6,8,10,12,15,20,25,30,35,40,45,50 bp) in a region of default size (200 bp) at the centre of each sequence. Following the Homer guidelines (http://homer.ucsd.edu/homer/index.html), the option -mask was used, to minimize the bias towards long repeats in the genome, and the maximum number of mismatches allowed in the global optimization phase has been set to 3, to improve the sensitivity of the algorithm. Other settings have been left as default, such as the distribution used to score motifs (binomial). Finally, a scoring algorithm assigned a ranked list of best matches (known motifs or genes) to each *de novo* motif, to inform the biological interpretation of the results.

### Image analysis

Whole-slide images (WSI) of immunostained sections of xenografts derived from GICs lines were analysed using QuPath^46^. Machine learning-based pixel classifiers were manually trained on a subset of WSI to detect tissue sections and vimentin immunostaining and create corresponding annotations. The trained pixel classifiers were then applied to the whole set of WSI. Prior to detecting vimentin immunostaining using the pixel classifier. Tissue annotations were eroded/shrunk by 35 μm to exclude tumours at the edge of the tissue sections.

Tumour core annotations were created by eroding/shrinking vimentin annotations by 10 μm to exclude any small and isolated staining as well as thin processes projecting from the tumour core. The annotation was then dilated/expanded to return the tumour core annotation to its original border size. Any annotations that were smaller than 10,000 μm^2^ were discarded to exclude small islands.

Gross tumour annotations were created by dilating/expanding vimentin annotations by 75 μm to fill in gaps between vimentin staining close together to be considered as part of the gross tumour. These were then eroded/shrunk by 100 μm and then dilated/expanded by 25 μm to smoothen and return to the original border size. Any gross tumour annotations that did not contain a tumour core annotation were discarded.

The tumour’s invasiveness index (II)^47^, independent of tumour size, was calculated as:

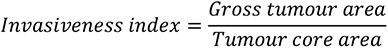

### Single cell RNA sequencing data analysis

Two public single cell RNA sequencing (scRNAseq) datasets have been analysed: tumour samples from seven newly diagnosed GBM patients (Antunes *et al.*^48^ patients ND1-ND7) and from eight GBM patients (Neftel *et al.*^8^ patients MGH102, MGH105, MGH114, MGH115, MGH118, MGH124, MGH125, MGH126, MGH143), both generated by the 10x Genomics platform. The gene expression matrices were downloaded from www.brainimmuneatlas.org, and GEO (accession number GSE131928), respectively. The gene expression matrices were merged, data was normalized, highly variable genes were detected, and their expression was scaled, followed by PCA, using the Seurat R package (Version 3.2.3). To account for the batch effect between samples, the cellular PCA embedding values were corrected with the harmony R package (Version 1.0), using a diversity clustering penalty parameter (theta) of one. Theta controls the level of alignment between batches, with higher values resulting in stronger correction. Next, the first 20 harmony corrected PCA embeddings were included in Louvain clustering (resolution = 0.25) and UMAP dimensionality reduction, using Seurat. The identified clusters were annotated as myeloid cells, lymphocytes, endothelial cells, and oligodendrocytes, based on expression of cell type markers. The remaining group of clusters were annotated as cancer cells.

The lymphocyte cluster was disaggregated and re-clustered, using 10 harmony corrected PCA embeddings and resolution=1. By using specific gene markers, the clusters were classified into B cells, Regulatory T cells, Proliferating CD8 T cells, NKT cells, Naive T cells, Interferon-response T cells, 2 clusters of CD4 T cells, 2 clusters of CD8 T cells and 2 clusters of NK cells.

The myeloid cells were also disaggregated and re-clustered based on 30 harmony corrected PCA embeddings, and resolution = 1. We identified Monocyte cluster, DC cluster, three clusters of microglia-derived TAMs (mg-TAMs), SEPP1-hi monocyte-derived TAMs (moTAMs), hypoxic moTAMs, IFN-response moTAMs, proliferating moTAMs, TAMs upregulating heat shock protein genes (HSP TAM: HSPA1A, HSPA1B, HSP90AA1, HSPH1, HSPB1), one cluster specific for patient MGH105, as well as myeloid-cancer cell doublets.

The cancer cells were disaggregated and re-clustered using 20 harmony corrected PCA embeddings, and resolution = 1. One of the clusters, which was expressing macrophage markers, was removed from further analysis as a cluster of macrophage-cancer cell doublets. Additive module scores for the astrocytic (ACS) and OPC gene signatures defined in this study were calculated for each cancer cell, using the AddModuleScore function of Seurat. This function yields the average expression level of each gene signature, subtracted by the average expression of a control gene set. Then, each cell has been assigned the signature with the highest score (ACS, OPC Enrich-300 or OPC Spec-300). Alternatively, additive module scores were calculated for the six gene signatures described by Neftel *et al.*^8^ (“MES1”, “MES2”, “AC”, “OPC”, “NPC1” and “NPC2”). The six gene signatures were defined using the genes in Supplementary TableS2 of Neftel *et al.*^8^. Each cell was also assigned the signature with the highest from the six scores. For the gene set enrichment pseudo-bulk analysis of the cancer cells, the raw UMI counts were summed for all cancer cells per gene for each patient using the aggregateAcrossCells function from the scuttle R package (Version 1.0.4). The pseudo-bulk counts were normalized using the CPM method and log2-transformation by edgeR (Version 3.32.1). Gene set ssGSEA enrichment scores for the ACS and OPC signatures, as well as for the six gene signatures of Neftel *et al.*^8^ were calculated using the GSVAR package (Version 1.38.2).

Spearman’s correlation coefficients have been calculated between ACS pseudo-bulk enrichment score divided by the mean between the two OPC scores per patient, and the percentages of the different cell populations per patient. Furthermore, Spearman’s correlation analysis was performed between the percentages of cells assigned to the distinct gene signatures per patient and the percentages of the different cell subsets present per patient. The data from patients ND7 and MGH143 was not used in these analyses, as they only contain CD45+, and CD45- sorted cells, respectively.

### Statistical analysis and graphs

All statistical analysis and generation of graphs was performed using GraphPad Prism 9 or R with appropriate R packages already mentioned. Parametric data are presented as mean ± standard deviation. P<0.05 was considered statistically significant, with p values <0.0332, <0.0021, <0.0002, <0.0001 represented with *, **, ***, **** respectively. Further information of the statistical analysis of specific datasets is indicated in the figure legends.

All scatter plots, time-series plots, bar graphs and survival curves, and accompanying statistical tests were generated with GraphPad Prism 9 or R. Venn diagrams were produced using the R package VennDiagram^49^, for the comparison and visualisation of gene lists. Fisher’s exact tests, used to test the significance of the overlap between Venn diagram categories, were calculated using an online statistics tool (https://www.socscistatistics.com/tests/fisher/default2.aspx) (no reference available). Upset plots showing the number of DEGs, found in various patient comparisons and the overlaps between patient comparisons were generated using the online tool Intervene Shiny App (https://asntech.shinyapps.io/intervene/) (no reference available).

## Data availability

Previously generated GIC and iNSC transcriptomic and methylation data are available from the NCBI Gene Expression Omnibus (GEO) database: GSE154958 and GSE155985. Newly generated transcriptomic and methylation data for iAPC, iOPC, iNSC and GIC are available as a private GEO submissions (GSE196418) (GSE196339) - please see the Transparent Reporting Form for this submission for instructions on how reviewers can access these data. The publicly available single cell dataset used in this study is available from the NCBI GEO database: GSE163120 and the European Genome-Phenome Archive: EGAS00001004871. GIC transcriptomic data from the HGCC are available from the NCBI GEO database: GSE152160, DNA methylation data from the HGCC are available from: http://portal.hgcc.se/data/HGCC_DNA_methylation.txt.

## Results

### A DNA hypo-methylation bias in a subset of GBM

To investigate the DNA methylome of GICs as compared to syngeneic iNSCs, we took advantage of a cohort of 10 previously described GIC/iNSC pairs^22^. DNA methylation was assessed using the Illumina Infinium Methylation EPIC array and processed as described in the Methods. Across the 10 intra-patient comparisons we visualised the distribution of the median probe delta M values and the proportion of these probes that were either hypo- or hyper-methylated, as well as the total number of differentially methylated regions (DMRs) and the proportion of hypo- and hyper-methylated DMRs (Figure 1 A, left panel). When comparing the proportion of hypo-methylated and hyper-methylated probes and DMRs for each comparison, four GICs stood out (19, 30, 31 and 17 – highlighted) identified as “bias” which had a statistically significant larger proportion of hypo-methylated probes and DMRs as compared to the other six GICs in the cohort, identified as “non-bias” (Figure 1 A, left panel and Figure 1 B). This subgrouping did not reflect the known DNA methylation-based classification^6^; three of the four GICs with a bias towards hypo-methylated DMRs belonged to the RTK-I (Proneural) subgroup and one to the RTK-II (Classical) subgroup. The GICs without a hypo-methylation bias were spread across the RTK-I, RTK-II and Mesenchymal subgroups. Noticeably, the proportion of hypo-methylated and hyper- methylated DMRs was exaggerated with a further increase in the proportion of hypo-methylated and hyper-methylated DMRs and probes respectively, when patient-specific probes and DMRs were considered (Figure 1 A, right panel).

**Figure 1:**
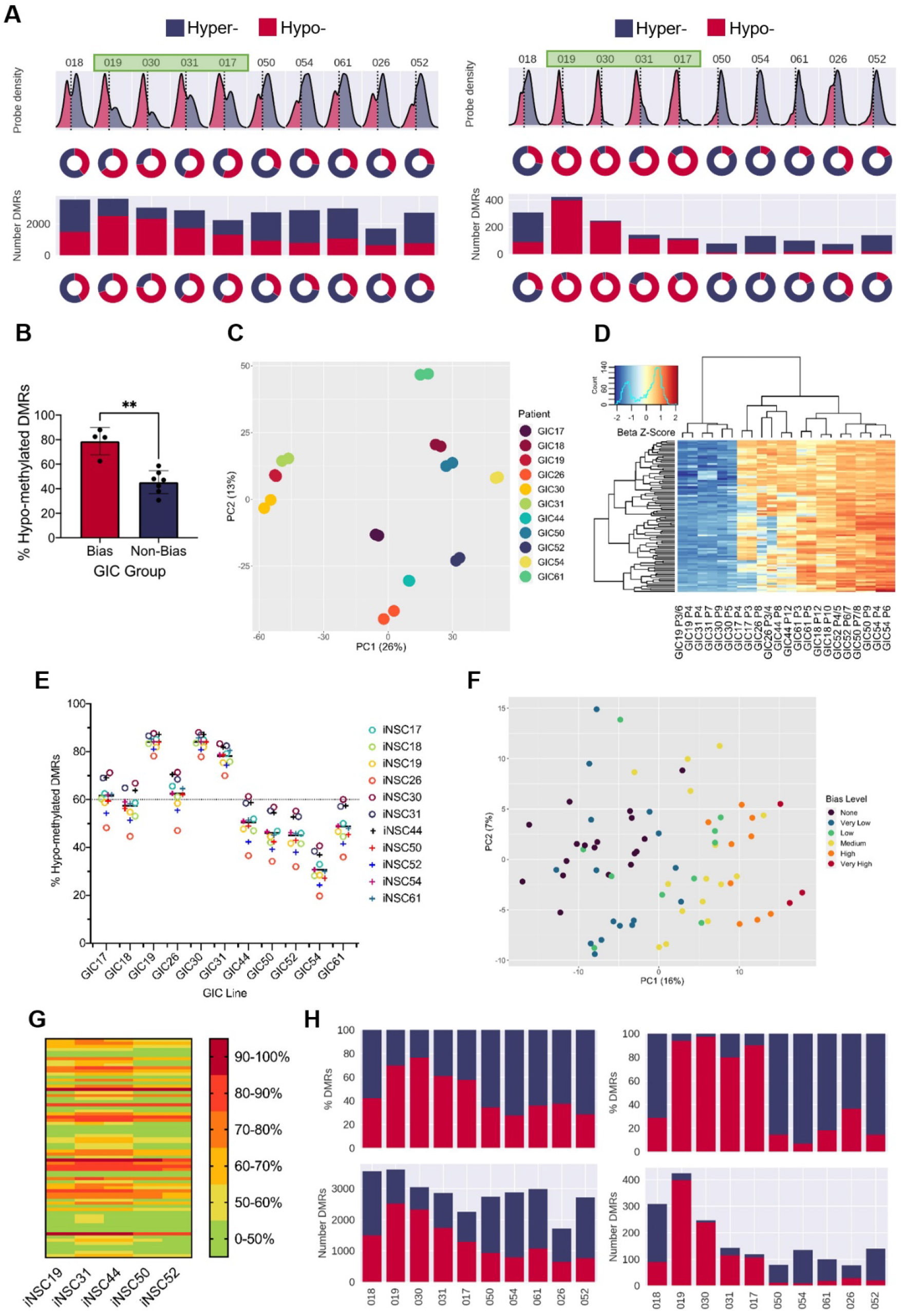
A DNA hypo-methylation bias in a subset of GBM. (**A**) The distribution of median probe delta M values (first row), the proportion of hypo- (red) and hyper- (blue) probes (second row), total number of DMRs (third row), and the proportion of hypo- and hyper- DMRs (fourth row) for each GIC - iNSC comparison (left panel) and using patient-specific probes and DMRs (right panel). (**B**) Percentage of hypo-methylated DMRs in bias GICs and non-bias GICs, statistical significance tested using Welch’s T-Test. (**C**) PCA of all patient-derived GIC samples from our cohort, based on the top 5000 most variable methylation probes. (**D**) Heatmap-dendrogram of the beta value z-scores of the top 100 methylation probes driving the variation of PC1 across all GICs and replicates in our cohort. (**E**) Percentage of hypo-methylated DMRs from all possible comparisons of GICs and iNSCs in our cohort, the mean percentage hypo-methylated DMRs for each GIC is represented by a horizontal black line. (**F**) PCA of all patient-derived GIC samples from the HGCC cohort, based on the top 5000 most variable methylation probes. (**G**) Heatmap summary of the percentage of hypo-methylated DMRs from all possible comparisons of HGCC GICs and 5 iNSCs from our cohort. (**H**) Number (bottom panel) and proportion (top panel) of hypo- and hyper- DMRs for each patient comparison between FFPE bulk tumour and iNSC, repeated in the right panel for patient-specific DMRs from the same comparisons.

Principal Component Analysis (PCA) of the 5000 most variable DNA methylation probes across all GICs in our cohort showed that Principal Component 1 (PC1) largely separated GICs by hypo-methylation bias, with three out of four GICs (19, 30 and 31) with a bias towards hypo-methylated DMRs clustering together on the left of the plot and all remaining patients to the right (Figure 1 C). A heatmap dendrogram of the beta values of the top 100 probes driving PC1 showed that these probes had much lower beta values in GICs 19, 30 and 31 relative to all other GICs, reflecting the separate cluster observed in the PCA (Figure 1 D). This observation suggested that the bias was GIC-driven, and not caused by the iNSC comparator, a conclusion which was confirmed when we performed non-syngeneic comparisons between each GIC and each of the iNSCs in our cohort (Figure 1 E), GIC17 was found to be an exception as the proportion of hypo-methylated DMRs was not as high as the other three GICs (19, 30 and 31), and indeed this GIC did not cluster with GICs 19, 30 and 31 in the PCA plot (Figure 1 C). However, the average percentage of hypo-methylated DMRs for this GIC, was still greater than 60% regardless of the comparator used. Interestingly, the average percentage of hypo-methylated DMRs for GIC26 was also greater than 60%, despite the cell line not meeting this threshold in the syngeneic comparison, possibly due to significant variability of the two biological replicates (Figure S1 J). The proportion of hypo-methylated DMRs for the remaining GICs varied from 20-70%, however the mean percentage of hypo-methylated DMRs was below threshold.

To validate this observation in an independent GIC cohort, DNA methylation data from the publicly available HGCC resource^50^ was used. The HGCC dataset contains 71 GIC samples, which were compared with iNSCs from our cohort, given that non-syngeneic comparisons do not prevent the identification of the hypo-methylation bias (Figure 1 E). Differential methylation analysis was performed between each HGCC patient-derived GIC line with one of our iNSC lines, and then repeated for five different iNSC lines to determine whether the results were consistent across comparisons. For each HGCC GIC, the mean percentage of hypo-methylated DMRs when compared against all iNSCs was then determined, to assess the extent of hypo-methylation bias for that GIC. HGCC GICs were stratified as either having no hypo-methylation bias (<50% hypo-methylated DMRs), a very low bias (>50%), low bias (>60%), medium bias (>70%), high bias (>80%) or very high bias (>90%). PCA of the 5000 most variable methylation probes across the HGCC GICs showed that PC1 separated patients on the extent of their hypo-methylation bias with very high bias GICs clustering to the right, and those with no bias clustering to the left (Figure 1 F). The reported percentage of hypo-methylated DMRs was found to be consistent regardless of the iNSC comparator used (Figure 1 G), in keeping with the results of the non-syngeneic comparisons performed in our own cohort (Figure 1 E). In total, 46.5% of HGCC GICs were found to have a hypo-methylation bias, comparable with the proportion in our smaller cohort (4/11, 36.4%). TCGA subgroup classification confirmed no enrichment for a specific subgroup (of the 33 GICs deemed to have a hypo-methylation bias, 15 were Mesenchymal, 4 Proneural, 6 Classical and 8 Neural), in keeping with the interpretation that the hypo-methylation bias observed is spread across the known GBM subgroups.

Finally, formalin fixed paraffin embedded (FFPE) tumour tissue, which was available for all 10 GIC/iNSC pairs of our cohort, was used as the neoplastic comparator to exclude that the observed hypo- methylation bias was induced in GIC by in-vitro culture. Once again, tumours 19, 30, 31 and 17 showed a greater proportion of hypo-methylated DMRs than the remaining six comparisons (Figure 1 H, left panel), and the proportion of hypo-methylated DMRs increased when only considering patient-specific DMRs (Figure 1 H, right panel).

In conclusion, we have identified a subset of glioblastoma that harbour a DNA hypo-methylation bias when either the bulk tumour or GIC derived thereof are compared to iPSC-derived NSC, herein referred to as hypo-bias GICs.

### The hypo-methylation bias does not globally impact on transcription

We used matched transcriptomic data from our GICs to assess any impact on transcription of the hypo- methylation bias. Unsupervised hierarchical clustering based on the 5000 most variably expressed genes was performed and did not recapitulate the grouping of GICs into the hypo-methylation bias group and non-bias group (Figure S1 A). Three out of four of the hypo-bias GICs (17, 19 and 30) grouped together along with one biological replicate from GIC26 – a line found to potentially harbour a hypo-methylation bias when non-syngeneic comparisons were performed, but not when syngeneic comparisons were performed. This grouping of GICs 17, 19, 26 and 30 did not correspond to their transcriptional GBM subgroup as they are RTKII, RTKI, MES and RTKI respectively. The remaining non-bias GICs largely clustered together along with one of the hypo-bias GICs – GIC31. Notably GIC44 clustered separately from all other patients. Furthermore, differential gene expression analysis between GICs and iNSCs did not reveal a bias in the directionality of differentially expressed genes (DEGs) when looking at all DEGs (Figure S1 B) or bias/non-bias group specific genes that are both differential expressed and differentially methylated (DE-DM) (Figure S1 C, D, E & F). Known genes associated with epigenetic remodelling or DNA methylation such as DNMTs or TETs were not identified within the DEGs (Supplementary File 1).

Next, we queried whether the hypo-methylation bias impacted miRNA expression and carried out small RNAseq. Clustering of the 500 most variably expressed miRNAs did not recapitulate the grouping of samples into hypo-bias and non-bias GICs (Figure S1 G). Similarly, comparisons between GICs and iNSCs did not reveal a bias in directionality of expression of differentially expressed miRNAs (Figure S1 H). Interestingly though, we identified five differentially expressed miRNAs common to hypo-bias GICs, that are differentially expressed in the same direction in all hypo-bias GICs (Figure S1 I). Some of these five differentially expressed miRNAs have previously been linked either directly or indirectly to neural development and/or glial lineage specification. Silencing of miR-1275 induces *GFAP* (an astrocyte marker) expression in glioblastoma cells^51^, and its down-regulation was associated with oligodendroglial differentiation of tumour cells^52^ – similarly in our dataset this miRNA is down-regulated. The *JAK*/*STAT* pathway, which is known to be a key regulator of astrocyte differentiation and activation^53–56^, is thought to be up-regulated by miR-4443, which is down-regulated in our GICs. Finally, miR-196, which is up-regulated in hypo-bias GICs, plays an essential role in neural development as it helps regulate Homeobox (*HOX*) genes^57^, some of which have been shown to play a role in astrocyte biology^58^.

In summary, the hypo-methylation bias identified here did not directly impact transcription or miRNA expression at global level, although differential expression of miRNAs involved in glial lineage specification was noted.

### Binding of transcription factors linked to DNA methylation and glial lineage specification are enriched at hypo-methylated loci in hypo-bias GICs

We used Homer^45^ – a tool to identify transcription factor binding sites that are enriched in a set of genomic loci-, to assess whether hypo-methylated DMRs from hypo-bias GICs (17, 19, 30 and 31) were enriched for specific transcription factor (TF) binding motifs, potentially linked to DNA methylation or glial lineage specification. Firstly, the hypo-methylated DMRs from our hypo-bias GICs, identified when comparing to syngeneic iNSCs, were compared against all other DMRs from all GIC – iNSC comparisons in our cohort. This showed that the top five enriched motifs from this comparison were matches for an array of Zinc-Finger proteins, *HOX* genes and families of factors such as Hepatocyte Nuclear Factor 4 (*HNF4*), Kruppel-Like Factors (*KLF*), Nuclear Factor I (*NFI*) and Distal-Less Homeobox (*DLX*) (Figure 2 A). We noticed that some of these TFs, namely *NFI*, *ETV4*, *DLX and KLF* have been linked to astrocyte/glial differentiation^59–61^.

**Figure 2:**
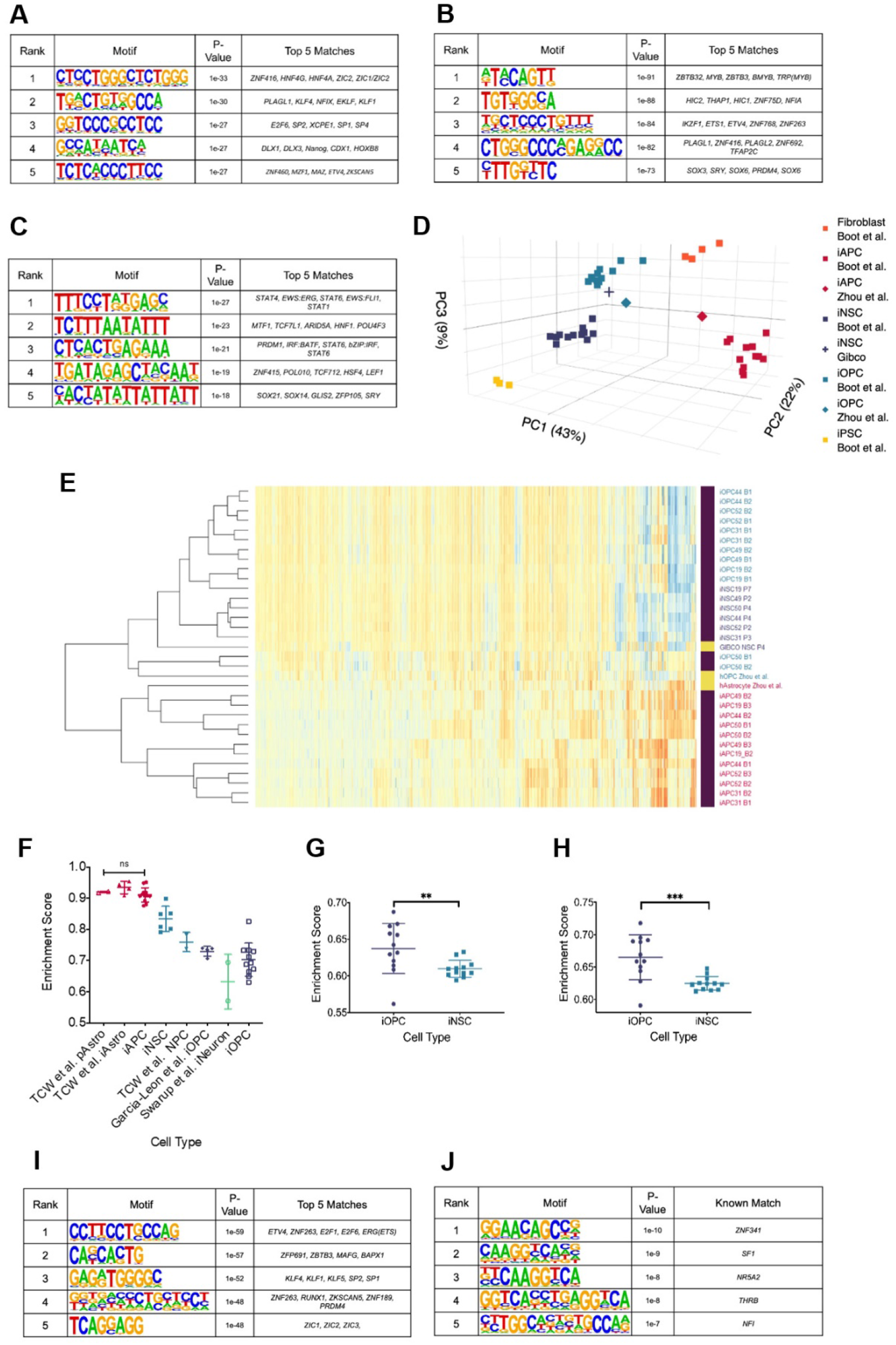
Hypo-methylated DMRs from hypo-bias GICs are enriched for transcription factor binding sites linked to glial differentiation. (**A**) Top five motifs enriched in all hypo-methylated DMRs from GICs 17, 19, 30 and 31 as compared to all other DMRs from all GICs. (**B**) Top five motifs enriched in all hypo-methylated DMRs from GICs 17, 19, 30 and 31 as compared to all the hypo-methylated DMRs from all other GICs. (**C**) Top five motifs enriched in all hyper-methylated DMRs from bias GICs as compared to all other DMRs from all GICs. (**D**) 3D PCA of methylation data from patient-derived fibroblasts, iPSCs, iNSCs, iAPCs, iOPCs, and publicly available reference datasets of NSCs, astrocytes and oligodendrocyte precursor cells. (**E**) Unsupervised hierarchical clustering based on the top 5000 variable methylation probes of iAPCs, iNSCs, iOPCs and publicly available reference datasets. (**F**) ssGSEA enrichment scores for the Astrocyte Composite Signature (ACS) of iAPCs, iNSCs, iOPCs and publicly available reference datasets, statistical differences tested with one-way ANOVA. ssGSEA enrichment scores of iOPCs and iNSCs for the Oligodendrocyte Specific-300 (**G**) and Oligodendrocyte Enriched-300 (**H**) gene signatures, statistical significance tested using Mann Whitney T-Test. Top five de-novo (**I**) and known (**J**) motifs enriched in all hypo-methylated DMRs in iAPCs, from each iAPC versus iNSC comparison.

To address the link to DNA methylation, Homer was used to identify the most enriched motifs in the hypo-methylated DMRs from the hypo-bias GICs, identified when comparing to syngeneic iNSCs, as compared to the hypo-methylated DMRs from the non-bias GIC comparisons. We reasoned that the hypo-methylated DMRs present in non-bias GICs are a background of hypo-methylated DMRs. Therefore, comparing against this background should identify transcription factors that have potentially contributed to the hypo-methylation of DMRs specifically in the hypo-bias GICs. When this comparison was performed, the top five enriched motifs were found to be matches for some of the same transcription factors identified in the first analysis such as *PLAGL1*, *NFI* family members and ETS Variant Transcription Factor 4 (*ETV4*) (Figure 2 B). Interestingly, the fifth ranked enriched motif from this analysis was a strong match for members of the *SOX* family, known to be involved in cellular differentiation and neural development^62^.

Next, Homer was used to identify the most enriched motifs in hyper-methylated DMRs from bias GICs, identified when contrasted to syngeneic iNSCs, as compared to all other DMRs from all other GIC – iNSC pairs. Here it was reasoned that hyper-methylated DMRs from bias GICs should not be enriched for any factors that may have caused the hypo-bias. Therefore, any motifs found to be enriched in this comparison were disregarded from further investigation. This comparison identified enriched motifs that were matches for various zinc finger proteins, members of the *STAT* family and *SOX* family. However, *PLAGL1*, *NFI* or *ETV4* were not identified in this analysis (Figure 2 C).

Therefore, we hypothesised that transcription factors such as the *NFI* family could be responsible for the hypo-methylation bias, as they have been shown to demethylate specific loci^59^, including lineage defining genes.

To test this hypothesis, we leveraged the availability of patient-specific iNSCs to generate induced astrocyte progenitors (iAPCs) from five different iNSC lines (patients 19, 31, 44, 50 and 52). DNA methylome and transcriptome were analysed at a differentiation stage, where cells co-expressed the astrocytic lineage marker *CD44*^63^ and the neural progenitor marker *NESTIN*^64^, in keeping with a progenitor rather than mature astrocytic state (Figure S2 A & B). Acquisition of a pro-inflammatory response upon IL6 treatment^25^, not observed in iNSC, and brisk proliferative activity (Figure S2 C & D), confirmed the astrocytic commitment of the cells. Induced oligodendrocyte progenitors (iOPC) were also generated for comparative analysis (Figure S2 E & F). Inspection of the DNA methylation and RNAseq datasets obtained from these cells, confirmed that iAPCs and iOPCs were epigenetically and transcriptionally distinct from one another and from the cells from which they were derived (Figure 2 D & E). Furthermore, both cell types were found to be enriched for an astrocytic and oligodendrocyte signature respectively (Figure 2 F, G & H).

Differential methylation analysis was then carried out in a syngeneic fashion, between each of the five iAPCs and their matched iNSC samples. Hypo-methylated DMRs in all iAPC lines were identified and Homer analysis performed comparing these sequences against the sequences of all other DMRs. When considering the so-called “*de novo*” Homer results, the top five enriched motifs were matches for binding sites previously found to be enriched in the comparison of hypo-methylated DMRs from hypo-bias GICs against all other DMRs, and the comparison of hypo-methylated DMRs from hypo-bias GICs against hypo-methylated DMRs from non-bias GICs (Figure 2 A & B). In particular, the top enriched motif was a strong match for *ETV4*, which was previously shown to be enriched in hypo-methylated DMRs from hypo-bias GICs thus confirming the link between this factor and hypo-methylated astrocyte associated loci (Figure 2 I). The *NFI* binding motif was the fifth-ranked enriched motif among the “*known matches*” from Homer (Figure 2 J). *ETV4* and *NFI* binding sites were not identified as enriched in either the “*de novo*” or “*known matches*” when Homer analysis was performed to compare hypo-methylated DMRs from iOPCs, identified when comparing to iNSCs against all other DMRs from this comparison (Figure S2 G & H). Instead, hypo-methylated DMRs in iOPCs as compared to iNSCs were enriched for the binding sites of *ZIC* genes, *HOX* genes, *SOX* genes and an array of zinc finger proteins (Figure S2 G & H).

In conclusion, hypo-methylated DMRs from hypo-bias GICs are enriched for transcription factor binding sites that have been linked to glial differentiation. iAPC hypo-methylated loci are similarly enriched for *ETV4* and *NFI* transcription factor binding sites, as are hypo-methylated DMRs from hypo-bias GICs, a phenomenon, which is specific to the astrocytic lineage. These results raise the possibility that GICs with a hypo-methylation bias have undergone glial priming prior to, or during, neoplastic transformation.

### A positive correlation between DNA hypo-methylation and astrocyte signature enrichment

Next, we set out to assess whether GICs with a hypo-methylation bias were enriched for specific signatures of the glial lineage. We performed single sample Gene Set Enrichment Analysis (ssGSEA) on our cohort of GICs for an early radial glia (early-RG) signature^65^, a bespoke astrocytic signature, termed Astrocyte Composite Signature (ACS) and two oligodendrocyte signatures (termed OPC Enriched-300 and OPC Specific-300), see Materials and Methods for further details on these signatures. All GICs in the cohort scored highly for the early-RG signature with little variability between lines (Figure S3 A), likely reflecting the high degree of transcriptional similarity between GICs and NSCs. However, four out of four hypo-bias GICs (17, 19, 30 & 31), as well as GIC44, were found to have a statistically significant higher enrichment score for the ACS than the two OPC signatures (Figure 3 A). GICs 50 and 26 also showed significant difference between the enrichment scores for the ACS and OPC signatures, however with a greater margin of error, possibly indicating more variability between the two biological replicates for these samples. All other GICs in the cohort showed no statistically significant differences in enrichment scores for all three gene signatures (Figure 3 A).

**Figure 3:**
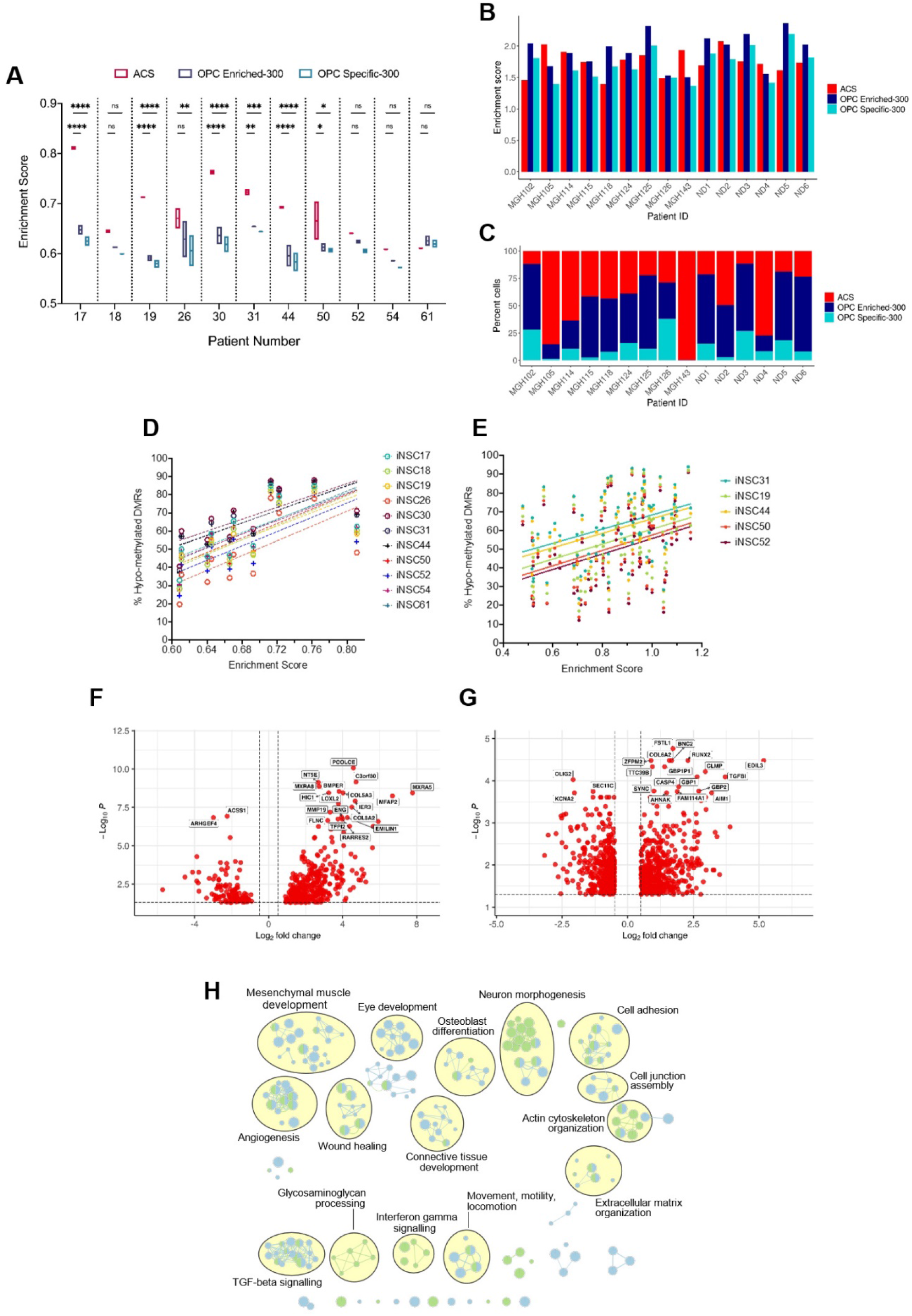
A positive correlation between DNA hypo-methylation and astrocyte signature enrichment. (**A**) ssGSEA enrichment scores of GICs (N = 2) from 11 GICs for three different gene signatures; ACS (red), OPC Enriched-300 Signature (blue) and OPC Specific-300 Signature (teal), statistical significance tested using two-way ANOVA. (**B**) ssGSEA enrichment scores based on pseudo-bulk (aggregated within a patient) data of the cancer cell subset from the scRNAseq glioblastoma tumour data from Antunes et al.^48^ and Neftel et al.^8^ for three different gene signatures, ACS (red), OPC Enriched-300 Signature (blue) and OPC Specific-300 Signature (teal). (**C**) Percentage of single cells for each tumour, which scored the highest for one of the three different signatures: ACS (red), OPC Enriched-300 Signature (blue) and OPC Specific-300 Signature (teal). (**D**) Scatter plot of the ACS enrichment score and the percentage of hypo-methylated DMRs for each of the GICs from our cohort, when comparing to each of the iNSCs from our cohort. (**E**) Scatter plot of the ACS enrichment score and the percentage of hypo-methylated DMRs for each of the HGCC GICs, when comparing to iNSCs from our cohort. (**F**) Volcano plot of DEGs identified from the comparison of bias/enriched GICs versus non-bias/non-enriched GICs (glm model used for DE analysis). (**G**) Volcano plot of DEGs identified from the comparison of bias/enriched GICs versus non-bias/non-enriched GICs from the HGCC cohort (glm model used for DE analysis). (**H**) Summary of pathway analysis performed using gProfiler, using the DEGs identified in (**F**) and (**G**).

To validate these results, ssGSEA was performed also on the GICs of the HGCC cohort^50^. A subset of GICs that are highly enriched for the ACS were identified also in this cohort (Figure S3 B). As there was only one GIC sample per patient present in the HGCC cohort, we determined a threshold for the enrichment scores that separated the samples into highly-enriched and lowly-enriched for the ACS signature. Firstly, an enriched cell line was defined as having a higher-than-average enrichment score for the ACS (samples had to score greater than 0.8429 for the ACS). Secondly, samples had to display an increased enrichment score as compared to the OPC signatures (ACS enrichment scores had to be at least 10% greater than any of the OPC signature scores). Given these two criteria, 55% of the samples were deemed to be enriched for the ACS in the HGCC dataset, hence confirming an ACS enrichment in an independent GIC cohort, compared to approximately 64% in our cohort.

Next, single-cell RNA sequencing (scRNAseq) data of glioblastomas from 2 different sources^8,48^ were interrogated to validate and further dissect, in glioblastoma tissue, the signature enrichment observed in a proportion of GIC. The analysis of the integrated scRNAseq dataset of 15 glioblastomas, showed presence of cancer cells, immune cells, oligodendrocytes, and a small cluster of endothelial cells. To identify patients enriched for the ACS signature in this dataset, we employed two strategies. First, by treating all the single cancer cells’ expression as pseudo-bulk tumour tissue data, we were able to identify a subset of tumours, which scored higher for the ACS than the two OPC signatures (MGH105, MGH114, MGH143, ND2 and ND4) (Figure 3 B). Alternatively, we determined enrichment scores for single cancer cells from each tumour and then assigned a signature identity to each cell to quantify the percentage of ACS, OPC-Enriched-300, or OPC-Specific-300 cells per tumour. Encouragingly the same tumours found to have a higher ACS enrichment score based on the pseudo-bulk analysis, had increased proportion of ACS cells: 75.07% ±19.40% compared to 26.26% ±11.69% in the remaining tumours (Figure 3 C). Finally, we applied these two approaches using the six cancer gene signatures from Neftel *et al.*^8^ (astrocytic: “AC”; mesenchymal: “MES1” and “MES2”; oligodendrocyte progenitor-specific: “OPC”; and neural progenitor-specific: “NPC1” and “NPC2”). The same five patients had higher enrichment scores and proportions of cancer cells, assigned to the astrocytic signature (“AC”) compared to the OPC signature, and the same held true for MGH115, MGH118, MGH124, MGH125, ND1 and ND6. For patients MGH143, ND2 and ND4, as well as for MGH115, MGH118 and ND5, the “AC” signature showed highest enrichment and percent assigned cells within the six studied gene modules (Figure S3 C & D). Interestingly, we noted that the ACS from our study enveloped the “MES1”, “MES2” and “AC” signatures from Neftel *et al.*^8^ (Figure S3 E & F) but scoring hypo-bias GICs from our cohort for the Neftel *et al.*^8^ signatures did not allow to further dissect the enrichment type as three out of four hypo-bias GICs (17, 19 and 30) scored most highly for the “MES1” and “MES2” signatures as well as the “AC” signature, GIC31 scored very similar for all Neftel *et al.* signatures^8^ (Figure S3 G).

A significant positive correlation (significantly non-zero) was found between the GIC ACS signature enrichment scores and the percentage of hypo-methylated DMRs in both our (Figure 3 D & S3 K) and in the HGCC cohorts (Figure 3 E & S3 L), raising the possibility that the two findings could be causally related.

To further explore the link between hypo-methylation bias and ACS enrichment, a differential expression analysis comparing hypo-bias GICs enriched for the ACS (termed bias/enriched), and GICs not harbouring a hypo-methylation bias and not enriched for the ACS (termed non-bias/non-enriched) was performed. For this analysis, GICs 17, 19, 30 and 31 were deemed to be bias/enriched, whilst GICs 18, 26, 50, 52, 54 and 61 were deemed to be non-bias/non-enriched. We reasoned that DEGs from this comparison could highlight key biological differences between bias/enriched and non-bias/non-enriched GICs. Interestingly, of the 465 DEGs (Figure 3 F), 47 were part of the ACS (Figure S3 H), – an overlap that was found to be statistically significant as tested by a Fisher’s exact test (p-value <0.00001). The same differential expression comparison in the HGCC cohort identified a total of 902 DEGs (Figure 3 G), of which a significant number (52) were also present in the ACS (Figure S3 I) (p-value <0.00001). When considering the top DEGs between the two groups in the HGCC cohort, *RUNX2* was up-regulated whilst *OLIG2* was down-regulated in bias/enriched GICs as compared to the non-bias/non-enriched GICs. This is an interesting finding, as *RUNX2* has been shown to drive astrocytic differentiation, whilst *OLIG2* is a master regulator of oligodendroglia differentiation^66^. An overlap of 105 genes from the 465 DEGs identified in our cohort and the 902 DEGs identified in the HGCC cohort was found, which was statistically significant as determined by Fisher’s exact test (Figure S3 J) (p-value <0.00001). Finally, pathway analysis on the two lists of DEGs, generated from our cohort and the HGCC cohort, revealed a shared enrichment for pathways associated with the extracellular matrix, morphogenesis, cell adhesion, angiogenesis, locomotion, wound healing, and cytokine signalling (Figure 3 H), suggesting a deregulation of these pathways in hypo-bias GICs, as compared to non-bias GICs.

In conclusion, we show in two independent GIC cohorts a positive correlation between the hypo-bias and the ACS enrichment score, in keeping with hypo-bias GICs being enriched for an astrocytic gene signature. An enrichment for an astrocytic signature is confirmed also in a proportion of glioblastoma at single cell level. Furthermore, when comparing bias/enriched GICs to non-bias/non-enriched GICs, a significant number of DEGs that are present in the ACS is found and a predicted impact on cell movement/invasion is identified.

### Increased invasion in xenografts derived from bias/enriched GICs and role of SRPX2 in regulating invasion in vitro

Next, we set out to assess whether bias/enriched GICs would give rise to tumours with distinct invasive properties as compared to those generated from non-bias/non-enriched GICs. Xenografts derived from our 10 GIC lines^22^ were stained for human vimentin on three levels. QuPath^46^, a machine learning-based pixel classifier was used for tissue detection from glass, vimentin staining detection and tumour core detection. We then calculated the invasiveness index (II) for each xenograft, defined as the ratio of area covered by infiltrating tumour cells to the area of the tumour core (gross tumour area/tumour core area) independent of tumour size^47^. Interestingly, xenografts derived from bias/enriched GICs showed increased invasiveness (Figure 4 A, B & C), despite having a smaller tumour core (Figure S4 A, B & C).

**Figure 4:**
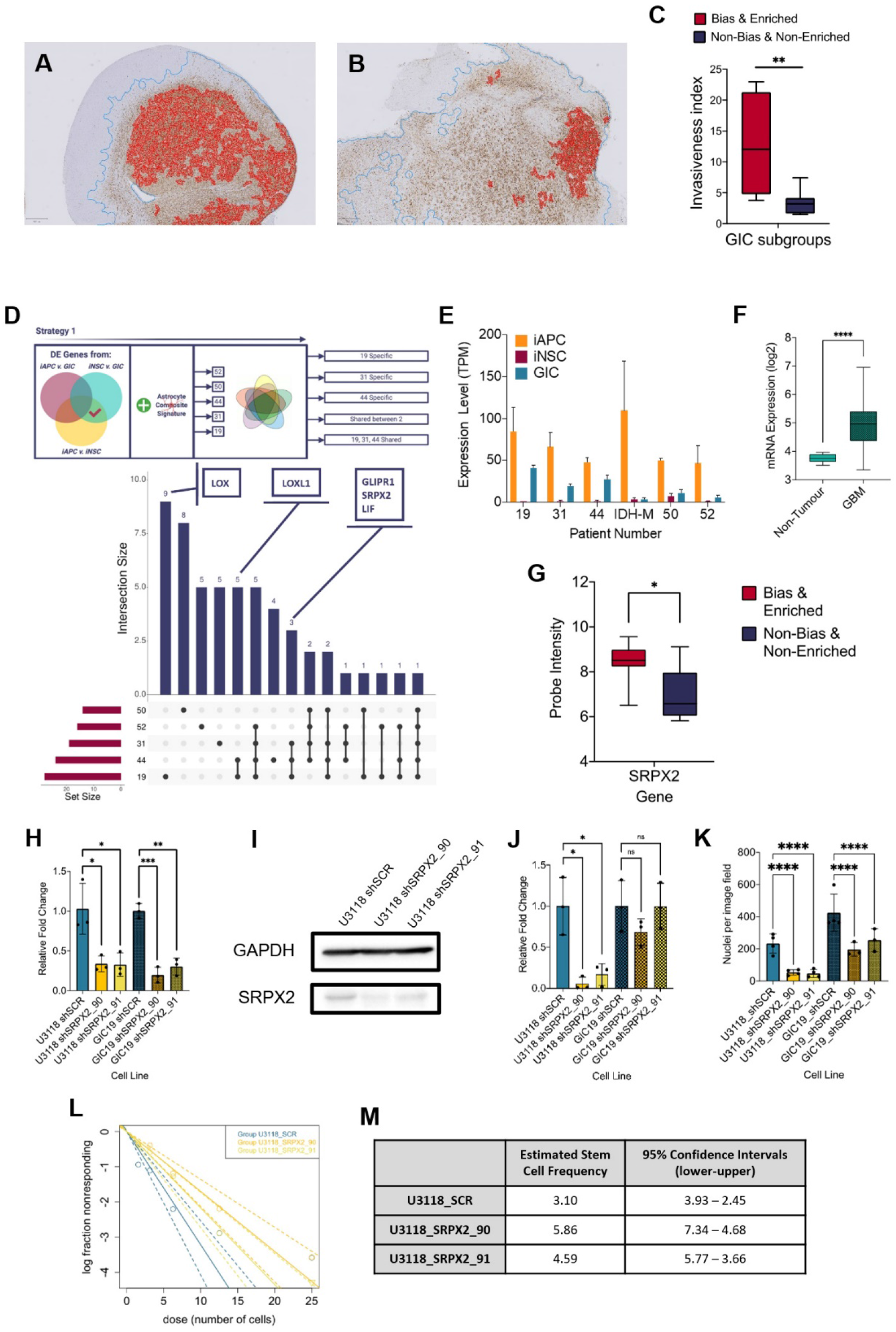
Increased invasion in xenografts derived from bias/enriched GICs and role of SRPX2 in regulating invasion in vitro. (**A** & **B**) Representative images of human vimentin-stained xenograft tumours with overlayed image analysis. Red outline is the detected tumour core, blue outline is the detected gross tumour edge. (**C**) Invasive index scores of xenografts from bias/enriched GICs and non-bias/non-enriched GICs, statistical significance tested using un-paired T-Test. (**D**) Overview of the strategy and upset plot of the number of DEGs identified in single or multiple patient comparisons. (**E**) Expression (TPM) of SRPX2 in the three cell types analysed: iAPC (orange), iNSC (red) and GIC (turquoise). (**F**) Expression of target genes in glioblastoma tissue as compared to non-tumour tissue, data acquired from Gliovis, statistical significance tested using Mann Whitney T-Test. (**G**) Expression of target genes in bias/enriched GICs vs non-bias/non-enriched GICs from the HGCC cohort, statistical significance tested using Mann Whitney T-Test. (**H**) Relative fold change in mRNA expression of SRPX2 as determined by qPCR for GIC shRNA knockdown lines, statistical significance tested using T-Test. (**I**) Representative western blot of SRPX2 in U3118 shRNA knockdown lines. (**J**) Relative fold change in SRPX2 protein expression as determined by western blot for GIC shRNA knockdown lines, statistical significance tested using T- Test. (**K**) Invasion assay results: average number of nuclei per image field of GIC SRPX2 knockdown lines, statistical significance tested using two-way ANOVA. (**L**) Neurosphere assay results: log fraction of the number of non-responding cultures at specified cell counts for U3118 SRPX2 knockdown lines. (**M**) Table of estimated stem cell frequencies and confidence intervals as determined by the neurosphere assay results and extreme limiting dilution assay analysis.

To identify genes that may contribute to the phenotypic characteristics of the bias/enriched GICs, and the tumours derived thereof, the following differential expression analyses were performed on a patient-by-patient basis: iAPC versus GIC, iAPC versus iNSC and iNSC versus GIC, with the aim to identify genes that play a role in the differentiation of iAPCs and are deregulated in GICs. Lists of DEGs for each patient comparison were then overlaid to find DEGs that were either unique or present in more than one comparison. DEGs present in both the iNSC versus GIC, and iNSC versus iAPC comparisons, were selected for further analysis as these DEGs could play a role in the differentiation of NSCs to astrocytes and because of the consequent similar expression level between GICs and iAPCs could indicate that GICs are ontogenetically linked to these progenitors.

We aimed at identifying genes that were differentially expressed in the two comparisons of interest and were at the same time part of the ACS (Figure 4 D). We focussed on DEGs that were specific to GICs 19, 31 and 44 and shared between all three. Three DEGs were shared between all three patient comparisons; these were GLI Pathogenesis Related 1 (*GLIPR1*), Sushi Repeat Containing Protein X-Linked 2 (*SRPX2*) and Leukaemia Inhibitory Factor (*LIF*) (Figure 4 D). Upon literature review *GLIPR1* was found to be very well studied in cancer^67^ and has been found to regulate migration and invasion of glioma cells^68,69^. Similarly, *LIF* is also well studied in glioma and has been shown to contribute to the maintenance of glioma-initiating cell self-renewal^70,71^. Moreover, *it* has been shown to help mediate astrocyte differentiation^72^. The pathogenic role and impact of *SRPX2* in GBM^73^ is less well characterised, despite been well studied in cancer^74–77^. Analysis of the pattern of expression of *SRPX2* in our data set found that it was up-regulated in the comparisons between iNSC and iAPC across all patients, and its expression was specifically higher in the GICs of patients 19, 31 and 44 (Figure 4 E), as expected. Further investigation found that *SRPX2* was more highly expressed in tumour tissue as compared to non-tumour brain tissue according to TCGA data (Figure 4 F) and in bias/enriched GICs from the HGCC cohort (Figure 4 G). To elucidate the role of *SRPX2*, the bias/enriched HGCC GIC line – U3118, and GIC19 were transduced with lentiviral vectors, carrying shRNA constructs targeting *SRPX2* (termed SRPX2_90 and SRPX2_91) and a scramble control. The mRNA levels of *SRPX2* were decreased upon silencing (Figure 4 H) and western blotting confirmed effective knockdown of the gene at the protein level (Figure 4 I & J & S4 D).

Because the comparison between bias/enriched and non-bias/non-enriched GICs identified cell movement/invasion among the differentially enriched pathways, a process which is also impacted in astrocytic progenitors, which are more motile as compared to NSC^78^ and because GIC of the bias/enriched subgroup gave rise to more invasive tumours upon intracerebral injection in mice, the impact of the SRPX2 knockdown on the invasive phenotype of the cells was assessed using a transwell invasion assay. The nuclei of cells that moved across the transwell membrane were fixed 24 hours after seeding, and then stained and counted as described in the Methods. On average, a statistically significant lower number of nuclei per image field were found in both GIC19 and U3118 *SRPX2* knockdown lines (Figure 4 K) (p-value <0.0001).

Proliferation was not affected in GICs upon *SRPX2* knockdown as compared to the scramble control lines (Figure S4 E) and variable results were obtained when self-renewal capacity was assessed by means of neurosphere extreme limiting dilution assay with less neurospheres forming in the U3118 shRNA *SRPX2* knockdown line at lower cell counts (6.25, 3.125 and 1.5625 cells per well) (Figure 4 L & M), but not in the GIC19 shRNA *SRPX2* knockdown lines (Figure S4 F & G).

In conclusion, we found that bias/enriched GICs gave rise to more invasive tumours in a xenograft model and that genes involved in migration/invasion are deregulated in these cells. Among these, *SRPX2* plays a role in essential properties of bias/enriched GIC, most notably in invasion, raising the possibility that *SRPX2* could be a therapeutic target for glioblastoma with a hypo-methylation bias and ACS enrichment.

### Glioblastoma enriched for an astrocytic signature display an altered immune landscape

Pathway enrichment analysis performed on the lists of DEGs, generated from the comparisons of bias/enriched GICs versus non-bias/non-enriched GICs identified immune-related pathways – TGF-beta signalling and interferon-gamma signalling – together with extracellular matrix organisation, locomotion, wound healing, morphogenesis, angiogenesis, and cell adhesion (Figure 3 H), raising the possibility of differences in the tumour microenvironment of these glioblastomas. Among the 47 DEGs shared with the ACS (Figure S3 I), Retinoic Acid Receptor Responder 2 (RARRES2 *aka.* Chemerin) – a chemoattractant, which binds to Chemerin chemokine-like receptor 1 (CMKLR1) - was noted. *RARRES2* was found to be significantly up-regulated in bias/enriched GICs compared to non-bias/non-enriched GICs, in both our own cohort and the HGCC cohort, and in non-G-CIMP tumour tissue relative to non-tumour brain tissue (Figure 5 A, B & Figure S5 A). Moreover, high expression of *RARRES2* is correlated with a worse prognosis (Figure S5 B). Significantly, *RARRES2* is well known to play a role in inflammation^79^ and promoting the migration of plasmacytoid dendritic cells, macrophages and NK-cells^80^ and its receptor CMKLR1, is expressed in tumour-associated macrophages (TAMs) from newly diagnosed GBMs (Figure 5 C & D), particularly in non-hypoxic TAMs (such as SEPP1-hi TAMs) as compared to hypoxic TAMs in recurrent glioblastomas (Figure 5 E & F).

**Figure 5:**
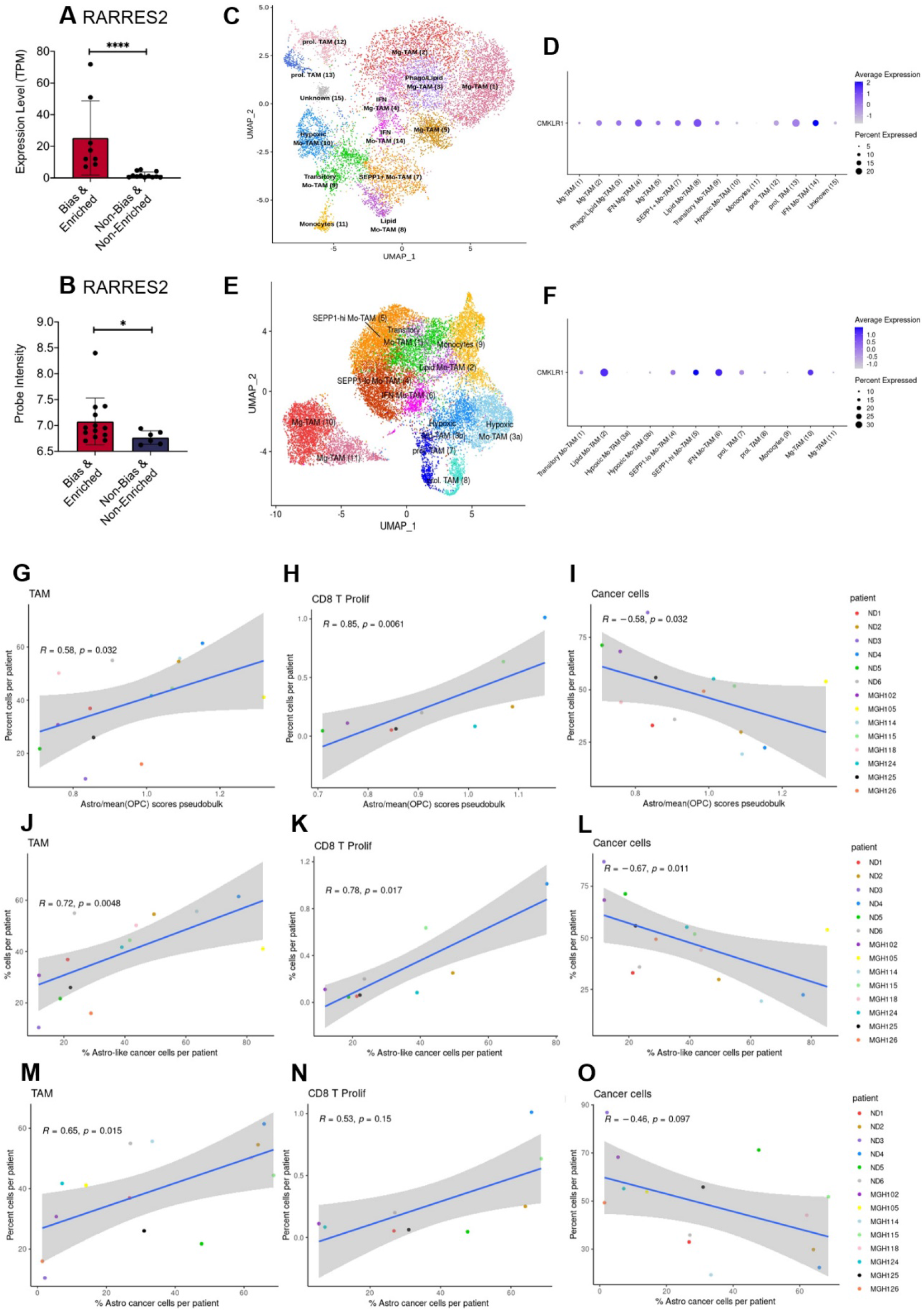
GBM enriched for an astrocytic signature display an altered immune landscape. (**A**) Expression of RARRES2 in bias/enriched GICs (N = 8) and non-bias/non-enriched GICs (N = 12) in our cohort, determined by RNAseq. (**B**) Expression of RARRES2 in bias/enriched GICs (N = 13) and non-bias/non-enriched GICs (N = 6) in the HGCC cohort. (**C**) UMAP of monocyte and TAM cell clusters from the Antunes et al.^39^ newly diagnosed glioblastoma tumour data; the cells are coloured by cell type. (**D**) Expression of CMKLR1 across different immune cell type clusters from panel (C). (**E**) UMAP of monocyte and TAM clusters from the Antunes et al.^39^, recurrent glioblastoma tumour data; the cells are coloured by TAM subtype. (**F**) Expression of CMKLR1 across different TAM subtype clusters from panel (**E**). Scatter plot, comparing the ratio of the ACS and the mean OPC pseudobulk enrichment scores, and the proportion of TAM cells (**G**), CD8 proliferative T-cells (**H**) and cancer cells (**I**) from the same tumour. Scatter plot, comparing the proportion of ACS-enriched cancer cells and the proportion of TAM cells (**J**), CD8 proliferative T-cells (**K**) and cancer cells (**L**) from the same tumour, corresponding to the dataset from (G - I). Scatter plot, comparing the proportion of AC-enriched and the proportion of TAM cells (**M**), CD8 proliferative T-cells (**N**) and cancer cells (**O**) from the same tumour, corresponding to the dataset from (G - I). Spearman’s rank correlation coefficient and the corresponding p-value are noted on each scatter plot. The blue lines represent smoothed conditional means using general linear model, while the grey areas on the plots denote the confidence interval around the smooth (using the geom_smooth function of ggplot2).

With this in mind, we leveraged the scRNAseq data from Antunes *et al.*^48^ and Neftel *et al.*^8^ to analyse the immune composition of tumours with an enrichment for an astrocytic signature or increased proportion of astrocyte-like cells (Figure 3 B & C). Both the myeloid and the lymphocyte clusters of the integrated scRNAseq data were selected and re-clustered, yielding various subpopulations of TAMs, dendritic cells (DC), mast cells, B, T, NKT and NK cells (Supplementary File 2 & 3). The ACS / OPC pseudo-bulk score ratio had a significant positive correlation with the proportions of TAMs (in particular of SEPP1-hi moTAMs and cDC2) and of proliferating CD8 T cells per patient (Figure 5 G & H). Conversely, a significantly negative correlation was found between the proportion of cancer cells and the ACS / OPC pseudo-bulk score ratio per patient (Figure 5 I). Similar trends were observed when, utilizing the single cell resolution of the scRNAseq data, we correlated the proportion of TAMs, proliferating CD8 T cells or cancer cells against the proportion of ACS-like tumour cells per patient (cells with higher ACS signature, compared to the two OPC signatures) (Figure 5 J – L). The comparison of the proportion of ACS-like cancer cells against the fractions of monocytes and IFN-response CD8 T cells per patient also yielded a significant positive correlation. Furthermore, we show that these results were reproduced, when the signatures from Neftel *et al.*^8^ were used to determine the proportions of astrocyte-like cancer cells per patient and the latter were correlated to the proportions of immune and cancer cells in each tumour (Figure 5 M – O). Finally, the opposite trends were observed when we correlated the proportion of TAMs, proliferating CD8 T cells or cancer cells against the proportion of OPC and NPC-like cancer cells (as determined by Neftel *et al.*^8^ signatures), however not all of the findings were statistically significant (Figure S5 C – H).

In conclusion, a significant up-regulation of *RARRES2* is identified in GICs with a hypo-methylation bias and ACS enrichment, and an increased proportion of TAMs and CD8 proliferative T-cells is found in tumours with a larger fraction of astrocyte-like tumour cells, indicating that the composition of the TME is different in tumours with an astrocytic signature enrichment.

## Discussion

We have identified a novel DNA hypo-methylation bias in a proportion of glioblastoma, which is ontogenetically linked to astrocyte progenitors. Hypo-methylated loci are enriched for binding motifs of transcription factors known to be involved in astrocyte differentiation, and differentially expressed genes and miRNAs known to play a role in astrocyte lineage commitment and differentiation were detected in hypo-bias GICs. At a functional level, bias/enriched GICs are characterised by increased invasion in vivo, enhanced *SRPX2*-regulated invasive properties in vitro, and an altered immune microenvironment.

Alterations of the DNA methylome have been previously described in genetically defined glioblastoma subtypes. In particular, the G-CIMP hyper-methylator phenotype characterises IDH-mutant gliomas^81^, where mutated IDH gains the ability to convert α-KG to 2-HG, which functions as an oncometabolite whilst also being a competitive inhibitor of α-KG dependent dioxygenases such as TET enzymes^81^. Consequently, IDH-mutant glioblastoma cells accumulate DNA methylation as TET enzymes are inhibited from removing methylation marks. A hypo-methylation bias has been previously found also in the paediatric H3 G34 mutant glioblastoma subgroup^6^, where mutations in the histone H3 variant H3.3 (H3F3A) block *SETD2* binding, leading to loss of H3K36 methylation, which in turn is linked to DNA methylation^82^. Alternatively, mutation and loss of *ATRX*, which is consistently found in G34 mutant glioblastoma could contribute to hypo-methylation at highly repeated sequences such as rDNA^83^. The DNA hypo-methylation bias described here is found in a proportion of IDH-wildtype glioblastomas, which are enriched for an astrocytic signature. The pan-cancer analysis of DNA methylomes, published in 2018^84^, reported that a proportion of IDH-wildtype gliomas exhibited high percentages of hypo- methylated loci, which correlated with a stemness signature^85^, based on hypo-methylation of specific loci enriched for the *SOX2-OCT4* binding motif. We did not find enrichment for the *SOX2-OCT4* motif in the hypo-methylated DMRs from hypo-bias GICs, but rather an enrichment for transcription factor binding sites known to play a role in glial/astrocyte differentiation, which is consistent with the enrichment for an astrocytic signature. Importantly, we have shown that the hypo-methylation bias is not induced in GIC by in-vitro culture as it was confirmed in the respective tumour tissue. Likewise, we have taken advantage of publicly available single cell transcriptome datasets to confirm that enrichment for our bespoke astrocytic signature as well as published astroglia signatures^8^ is found in a proportion of GBM.

Of particular interest was the finding of an enrichment for members of the *NFI* family in the hypo- methylated DMRs from hypo-bias GICs as previous studies have shown that astrocyte differentiation requires Nfia-induced demethylation of key astrocyte lineage specifying genes^59,86^. In fact, Sanosaka *et al.*^59^ showed that the methylome underpins the differentiation potential of NSCs rather than gene expression itself. They found that E11.5 embryonic mouse NSCs were lineage-restricted and only giving rise to neurons, whilst E18.5 NSCs had an increased proportion of hypo-methylated loci and were multipotent – being able to give rise to glial and neuronal lineages. Furthermore, this study also showed that DMRs with reduced methylation (hypo-methylated) in E18.5 NSCs as compared to E11.5 NSCs are enriched for *NFI* binding sites, and that this gene family was responsible for the loss of DNA methylation and gain of multipotency for the glial lineage. It is intriguing that these previous studies may provide an interpretative framework as to why we find a positive correlation between the hypo- methylation bias and ACS enrichment. It is conceivable that GICs with a hypo-methylation bias may arise from a neural progenitor population that has undergone priming for glial differentiation, for example by the *NFI* transcription factor family. Other transcription factor-binding sites we found enriched in hypo- methylated loci are also known to be involved in glial/astrocyte differentiation, such as *ETV4*^66,87^ *and PLAGL1*, the latter by transactivating *Socs3*, a potent inhibitor of pro-differentiative *Jak*/*Stat3* signalling, thereby preventing precocious astroglia differentiation^88^. We found a significant enrichment for genes of the ACS in the DE genes between hypo-bias and non-hypo bias GICs in both our and the HGCC datasets. Likewise, the 5 DE miRNAs identified in this comparison – miR-4443, miR-1275, miR-196, miR-5100 and miR-1268, which have been previously linked to cancer or glioblastoma pathogenesis^51,89–92^, have also been linked either directly or indirectly to glial differentiation. These include miR-196 known for its regulatory role of Homeobox (HOX) genes^57^ in both a healthy and malignant developmental context and miR-1275 previously shown to be involved in glial lineage specification^51–52^. Taken together, the results of the binding motif analysis and of the differential expression analysis raise the possibility that bias/enriched GICs arise from a neural progenitor, which has undergone glial/astrocyte priming. Overexpression of *NFI* family members in non-bias/non-enriched GICs will be required to assess whether changes in DNA methylation and gene signature enrichment can be elicited, or whether this enrichment is a consequence of the hypo-methylation bias. Alternatively, disruption of the *NFI* binding motifs across the genome, for example by means of CRISPR- Cas9 system^93^ could be carried out to assess the effect on DNA hypo-methylation and gene signature enrichment.

Importantly, when comparing genes differentially expressed between bias/enriched GICs to non-bias/non-enriched GICs, a predicted impact on cell movement/invasion was identified. In keeping with this prediction, xenografts generated from bias/enriched GICs were found to grow more invasively as compared to those generated by non-bias/non-enriched GICs. This is of particular interest in glioblastoma, given the diffusely infiltrative growth of these tumours, which plays a significant role in limiting the effectiveness of the current therapies. Among the genes deregulated, *SRPX2* stood out as it has been previously shown to be associated with poor prognosis, and to promote tumour progression and metastasis in primary GICs^73^. Indeed, we show that silencing of this gene leads to impaired invasion in two bias/enriched GIC lines in *in vitro* assays.

The immune microenvironment plays a crucial role in tumour pathogenesis, including glioblastoma^48,94^. We have identified a significant deregulation of genes involved in immunomodulatory pathways in bias/enriched GICs and analysis of scRNASeq datasets of glioblastoma has confirmed an altered immune landscape in glioblastoma with an ACS signature enrichment. The significant correlation between the ACS signature enrichment and the number of TAMs is of particular interest given the upregulation of *RARRES2* in bias/enriched GICs and the role of TAMs in glioblastoma invasion^95^. Interestingly, a SEPP1-hi phenotype was observed in these TAMs corresponding to an anti-inflammatory phenotype^48^, which raises the possibility that they could play a pro-tumourigenic role in this glioblastoma subgroup.

A recent methylome analysis of various tumour types has found that global methylation loss correlates with increased resistance to immunotherapy and immune evasion signatures^96^, hence the identification of this subgroup of GBM could have important implications in patient stratification for immunomodulatory treatments.

## Acknowledgements

This work is funded by grants from Brain Tumour Research (Centre of Excellence award to S.M.), Cancer Research UK (C23985/A29199 programme award to S.M.), Barts Charity (MGU0447 programme grant to S.M.). Part of the study was funded by the National Institute for Health Research to UCLH Biomedical research centre (BRC399/NS/RB/101410 to S.B.). S.B. is also supported by the Department of Health’s NIHR Biomedical Research Centre’s funding scheme. We acknowledge the use of data generated by the TCGA Research Network: https://www.cancer.gov/tcga.

## Author contribution

J.R.B. and S.M. designed experiments. J.R.B., X.Z. performed experiments and analysed results. G.R., N.P. and J.R.B. designed and performed all non-single cell computational analyses. S.B. obtained ethical approval and supervised human sample collection and pathological and molecular analysis of the biopsies. Y.M.L. performed image analysis. C.V., L.G. characterised GIC and iNSC lines and C.V. performed the intracranial-injection experiments. D.K. K.M. designed and performed all single cell computational analyses. M.R.B, D.S. and S.N. shared datasets and essential expertise. S.M. supervised the study and provided financial support. J.R.B. and S.M wrote the manuscript with contribution from all authors.

