## Supplementary Figures for "Global hypo-methylation in a subgroup of glioblastoma enriched for an astrocytic signature is associated with increased invasion and altered immune landscape"

***Supplementary Figures and Legends***

**Figure S1**

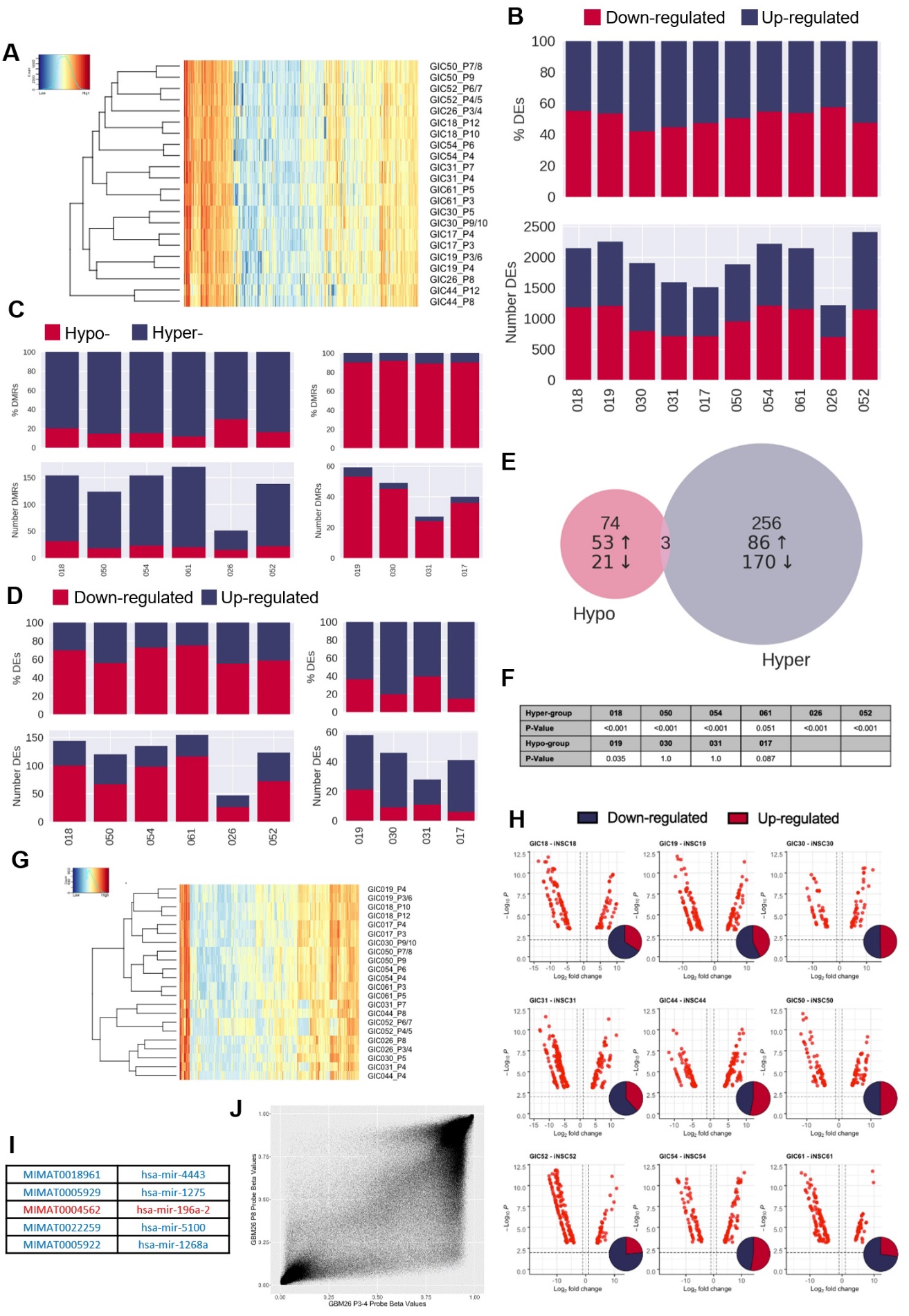

**Figure S1:** **The hypo-methylation bias does not globally impact on transcription.**

(**A**) Unsupervised hierarchical clustering of the top 5000 variably expressed genes across two biological replicates from the 11 GICs in our cohort. Euclidean clustering method and complete distance method used. Red indicates highly expressed, blue lowly expressed. (**B**) Total number (bottom panel) and proportion (top panel) of DEGs that are up-regulated (blue) or down-regulated (red) in GIC as compared to iNSC for each patient comparison. (**C**) Total number (bottom panel) and proportion (top panel) of DE-DM genes that are hyper- (blue) or hypo-methylated (red) in GIC as compared to iNSC for each patient comparison in the non-bias patient group (left) and the hypo-bias patient group (right). (**D**) Total number (bottom panel) and proportion (top panel) of DE-DM genes that are up- (blue) or down-regulated (red) in GICs as compared to iNSC for each patient comparison in the non-bias patient group (left) and the hypo-bias patient group (right). (**E**) Overlap in hyper- and hypo- specific DE-DM genes, and the direction of change in expression. (**F**) P-value summary of Fishers exact test performed to test the significance of the concordance between direction of DMRs and correspond DEGs. (**G**) Unsupervised hierarchical clustering of the top 500 variably expressed miRNAs across two biological replicates from the 11 GICs in our cohort. Euclidean clustering method and complete distance method were used. Red indicates highly expressed, blue lowly expressed. (**H**) Volcano plots of DE miRNAs (FDR <0.01, logFC >1) for each syngeneic comparison between iNSC and GIC with inset Venn diagram showing proportion of up-regulated (red) and down-regulated (blue) miRNAs. (**I**) List of five DE miRNAs common to patients 19, 30 and 31, blue indicating down-regulation in GIC, red indicating up-regulation. (**J**) Correlation of methylation probe beta values between the two technical replicates of GIC26.

**Figure S2**

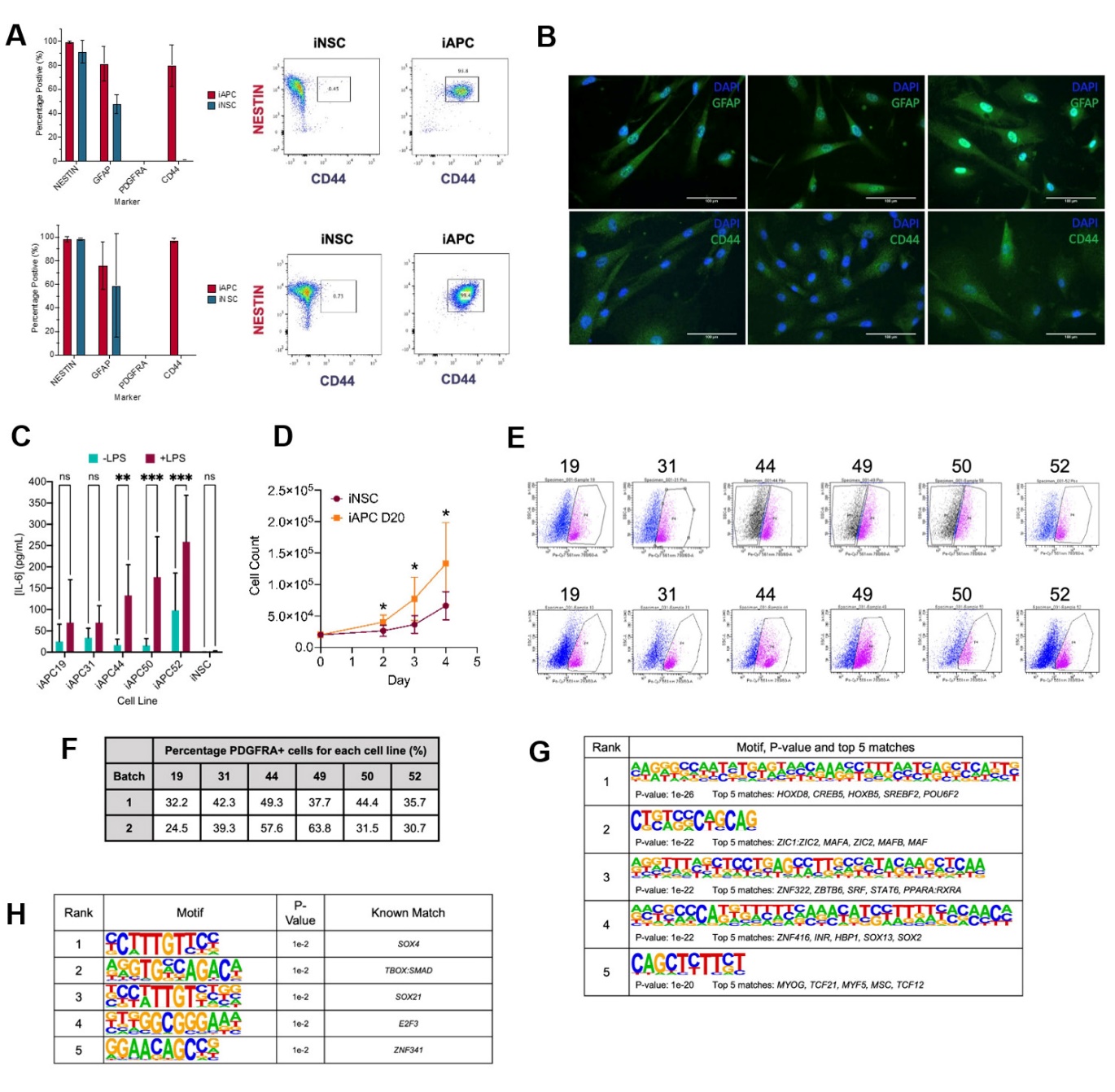

**Figure S2:** **Characterisation of iAPC and iOPC obtained from iNSC.**

(**A**) Flow cytometry results of iAPCs analysed after 10 days of differentiation (top panel) and 20 days (bottom panel). iAPCs (red) (N = 6) and iNSCs (teal) (N = 3) were analysed for the markers NESTIN, GFAP, PDGFRA and CD44. Results presented as bar charts of percentage of positive cells for respective markers, and for each time-point representative dot plots of NESTIN versus CD44 of iNSC and iAPCs, to show co-expression of the two markers. (**B**) Immunostaining of iAPCs, for GFAP (top panel, green) and CD44 (bottom panel, green) with DAPI nuclear staining (blue), after 30 days of differentiation. (**C**) Concentration of IL-6 produced by iAPCs and iNSCs stimulated with 50 µg of LPS for 24-hours (turquoise) and unstimulated (burgundy). Statistical significance was tested using a two-way ANOVA (N = 5 – 7). (**D**) Cell counts of iNSCs (maroon) and iAPCs differentiated for 20 days (iAPC D20) (orange). Cell counts at Day 2, 3 and 4 were statistically compared using a T-Test (N = 6). (**E**) FACS results, based on PDGFRA sorting, for Batch 1 (top panel) and Batch 2 (bottom panel) of iOPC differentiation. (**F**) FACS results summary: the percentage PDGFRA^+^ cells for each cell line from each batch. Top five de-novo (**G**) and known (**H**) motifs enriched in all hypo-methylated DMRs in iOPCs, from each iOPC versus iNSC comparison.

**Figure S3**

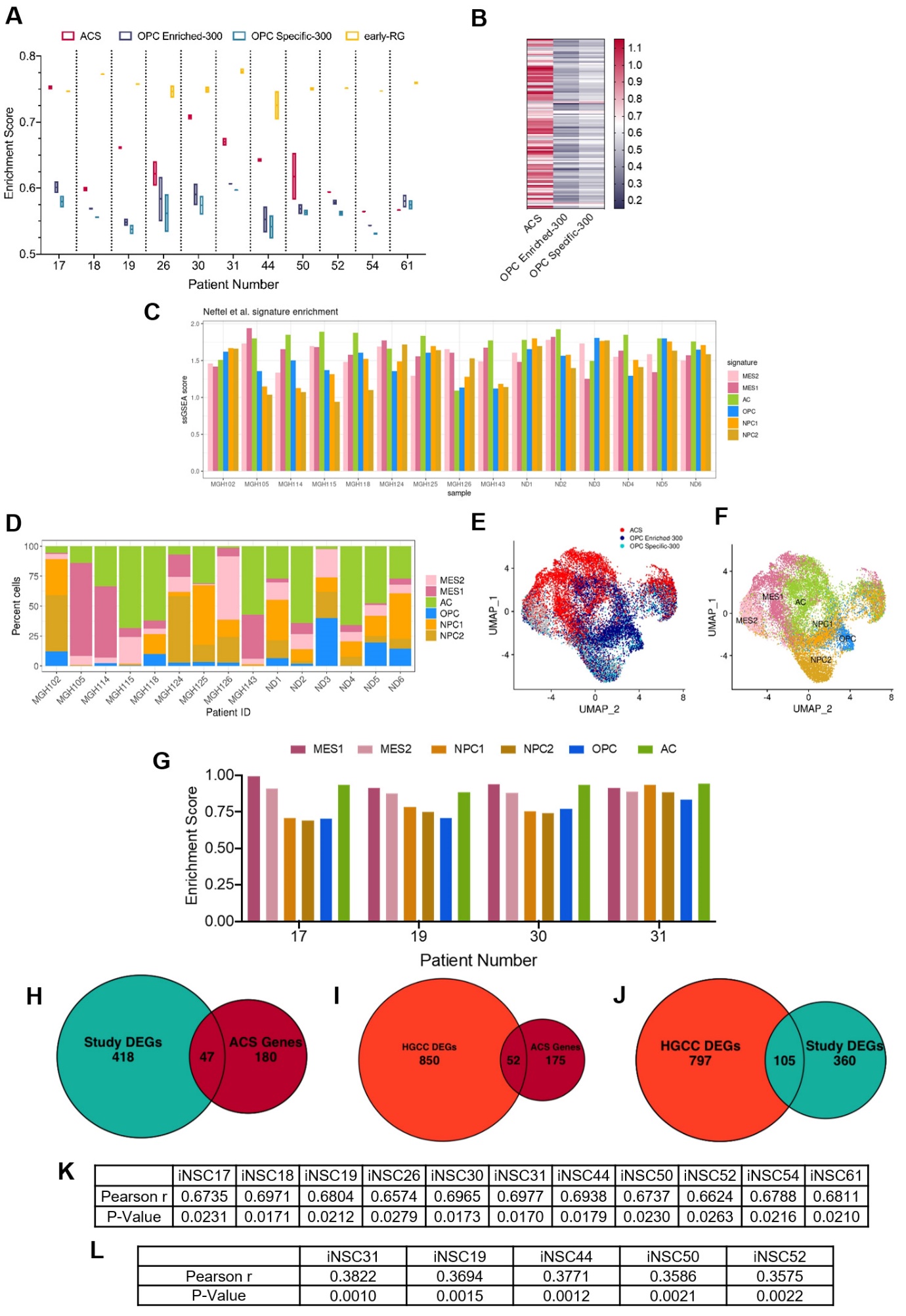

**Figure S3: Characterisation of the positive correlation between DNA hypo-methylation and astrocyte signature enrichment**

(**A**) ssGSEA enrichment scores of GICs (N = 2) from 11 GICs for four different gene signatures; ACS (red), OPC Enriched-300 Signature (blue) and OPC Specific-300 Signature (teal), and early-RG (yellow). (**B**) Enrichment score heatmap for the ACS, OPC Enriched-300 and OPC Specific-300 signatures, of external GICs (N = 1). (**C**) ssGSEA enrichment scores, based on pseudo-bulk (aggregated within a patient) data of the cancer cell subset from the scRNAseq GBM tumour data from Antunes et al.^1^ and Neftel et al.^2^ for six cancer gene signatures from Neftel et al.^2^ (**D**) Percentage of single cancer cells for each tumour, corresponding to the dataset from (**C**), which scored the highest for one of the six signatures from Neftel et al.^2^ (**E**) UMAP plot of single cancer cells, corresponding to the dataset from (**C**); each cell was assigned to the signature with the highest enrichment score between ACS, OPC Enriched-300 or OPC Specific-300. The cells were coloured based on the assigned signature type. (**F**) UMAP plot of single cancer cells, corresponding to the dataset from (**C**); each cell was assigned to the signature with the highest enrichment score from the Neftel et al.^2^ signatures. The cells were coloured based on the assigned signature type. (**G**) ssGSEA enrichment scores of GICs (N = 2) from 11 GICs for the six different signatures from Neftel et al.^2^ (**H**) Overlap between DEGs identified from the comparison of bias/enriched GICs versus non-bias/non-enriched GICs (from our cohort) and the ACS genes, Fisher exact test p-value < 0.00001. (**I**) Overlap between DEGs identified from the comparison of bias/enriched GICs versus non-bias/non-enriched GICs (from the HGCC cohort) and the ACS genes, Fisher exact test p-value < 0.00001. (**J**) Overlap between DEGs identified when comparing bias/enriched GICs versus non-bias/non-enriched GICs from our cohort and the HGCC cohort, Fisher exact test p-value <0.00001. (**K**) Pearson r- and P-values of correlations between percentage hypo-methylated DMRs (when compared to iNSCs) and ACS enrichment score for GICs from our cohort, corresponding to Figure 3 B. (**L**) Pearson r- and P-values of correlations between percentage hypo-methylated DMRs (when compared to iNSCs) and ACS enrichment score for GICs from the HGCC cohort, corresponding to Figure 3 C.

**Figure S4**

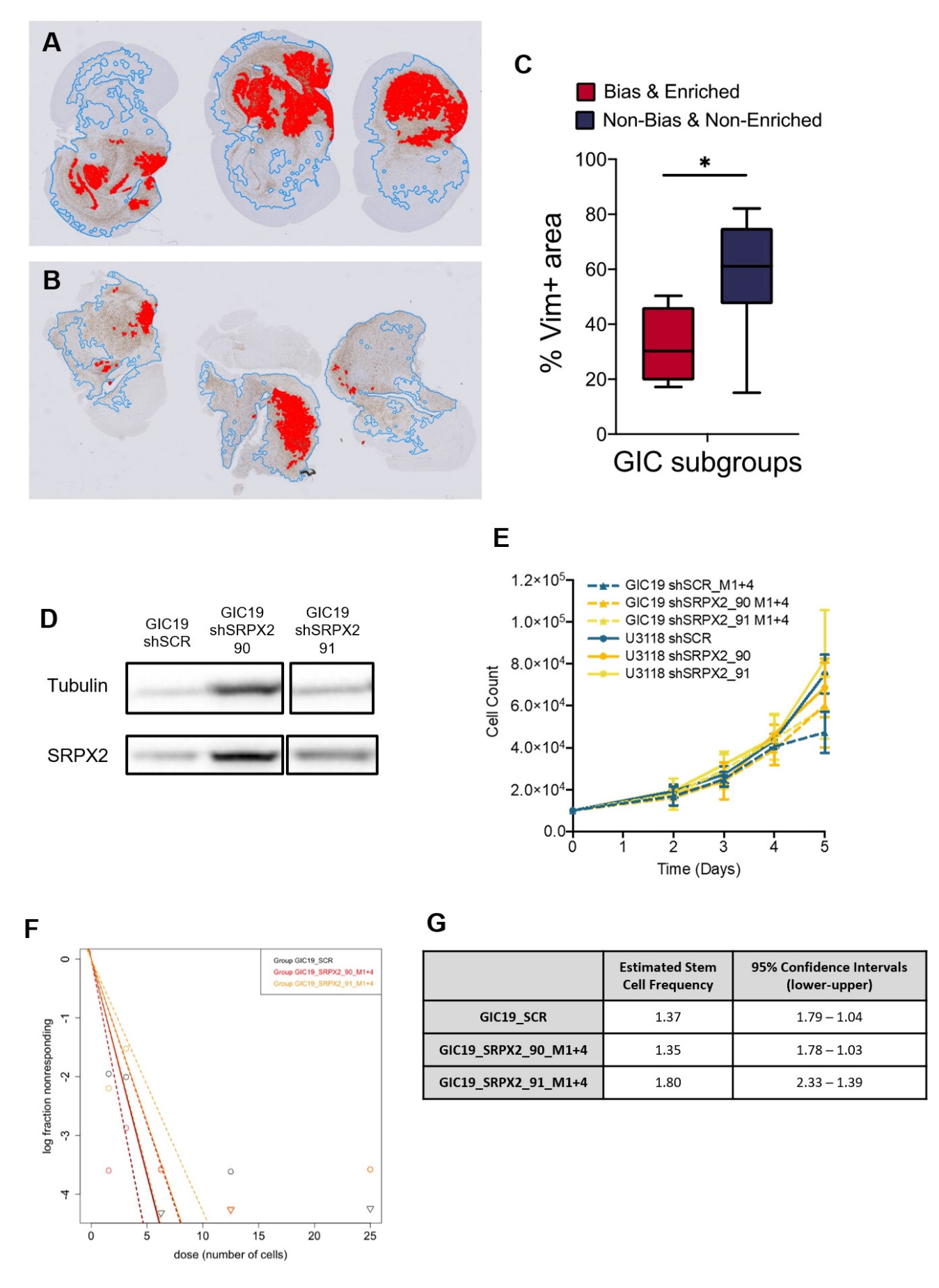

**Figure S4: Characterisation of xenografts derived from bias/enriched GICs and role of SRPX2 in regulating GIC properties.**

(**A** & **B**) Representative overview images of human vimentin-stained xenograft tumours with overlayed image analysis. Red outline is the detected tumour core, blue outline is the detected gross tumour edge. (**C**) Percentage vimentin positive area relative to tissue area of xenografts from bias/enriched GICs and non-bias/non-enriched GICs, statistical significance tested using un-paired T-Test. (**D**) Representative western blot of SRPX2 in GIC19 shRNA knockdown lines. (**E**) Proliferation assay growth curves for U3118 and GIC19 SRPX2 knockdown lines. (**F**) Neurosphere assay results: log fraction of the number of non-responding cultures at various cell counts for GIC19 SRPX2 knockdown lines. (**G**) Table of estimated stem cell frequencies and confidence intervals as determined by the neurosphere assay results and extreme limiting dilution assay analysis.

**Figure S5**

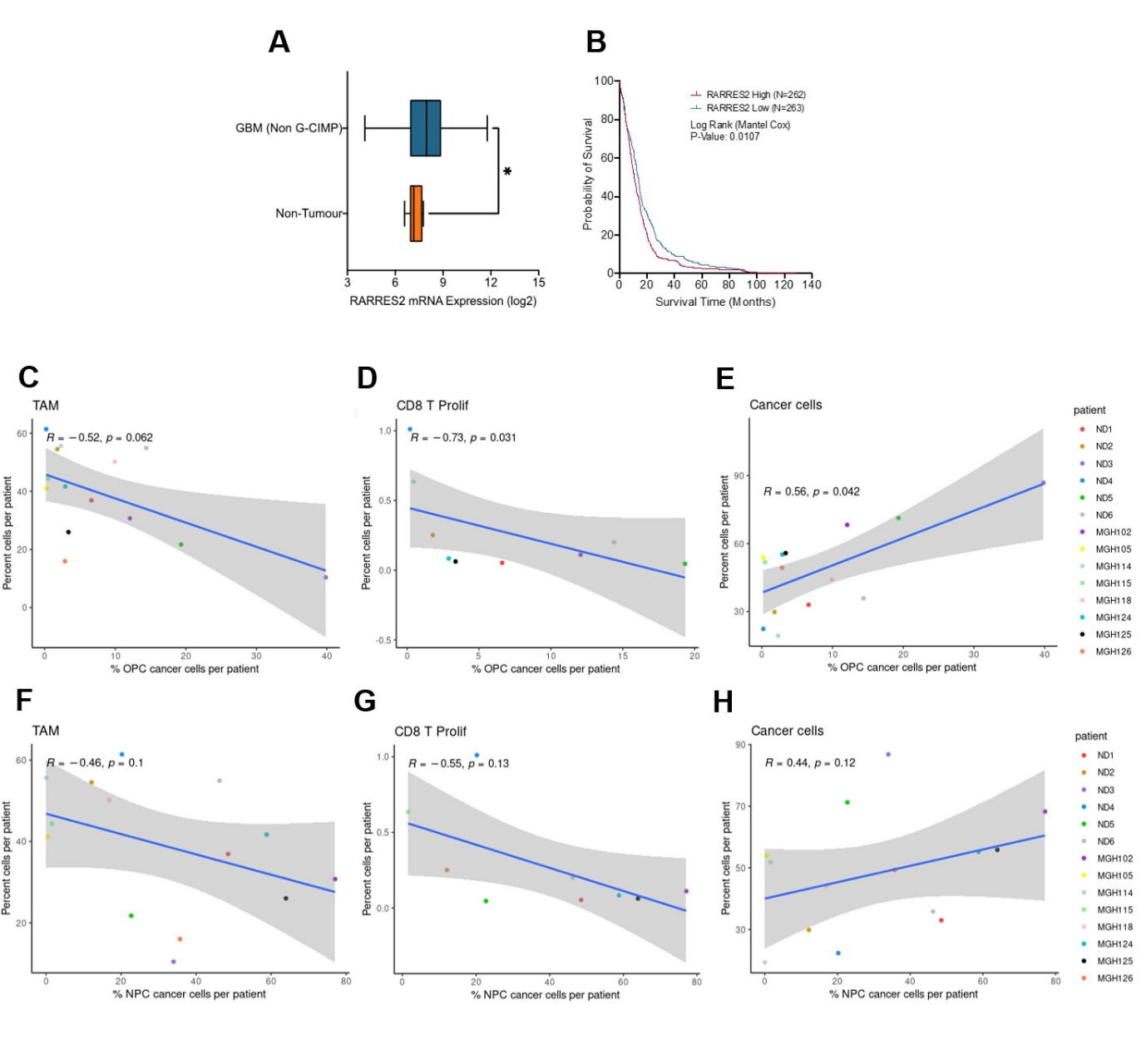

**Figure S5: scRNA analysis of the TME composition in glioblastoma enriched for an astrocytic signature**

(**A**) Expression of RARRES2 in non-G-CIMP tumour tissue (N = 482) as compared to healthy non-tumour tissue (N = 10), statistical significance tested using Mann Whitney T-Test, produced using TCGA data available on Gliovis. (**B**) Kaplan-Meier curve for GBM patients with high expression (red) versus low expression (blue) of RARRES2, produced using TCGA data available on Gliovis. Scatter plot, comparing the proportion of OPC-enriched cancer cells (as determined by the gene signatures from Neftel et al.^2^) and the proportion of myeloid cells (**C**), CD8 proliferative T-cells (**D**) and cancer cells (**E**) from the same tumour of the scRNAseq GBM tumour data from Antunes et al.^1^ and Neftel et al.^2^ Scatter plot, comparing the proportion of NPC enriched tumour cells (as determined by the signatures^2^) and the proportion of myeloid cells (**F**), CD8 proliferative T-cells (**G**) and cancer cells (**H**) from the same tumour, corresponding to the dataset from (C-E).
